# Nuclear lipidome is altered in amyotrophic lateral sclerosis: a preliminary study

**DOI:** 10.1101/682526

**Authors:** Omar Ramírez-Nuñez, Mariona Jové, Pascual Torres, Joaquim Sol, Laia Fontdevila, Ricardo Romero-Guevara, Victòria Ayala, Chiara Rossi, Jordi Boada, Mònica Povedano, Pol Andrés-Benito, Isidro Ferrer, Reinald Pamplona, Manuel Portero-Otin

**Affiliations:** Department of Experimental Medicine, School of Medicine, IRBLleida-UdL, Avda Rovira Roure 80 25196, Lleida, Spain; Neurology Service, Hospital Universitari de Bellvitge, L’Hospitalet de Llobregat, c/La Feixa Llarga, S/N 08908 Hospitalet de Llobregat, Barcelona, Spain; Departament of Pathology and Experimental Therapeutics, Universitat de Barcelona; Hospital Universitari de Bellvitge, IDIBELL, Hospitalet de Llobregat; CIBERNED (Centro de Investigación Biomédica en Red de Enfermedades Neurodegenerativas), Instituto Carlos III, c/La Feixa Llarga sn, 08908 Hospitalet de Llobregat, Barcelona, Spain

**Author notes:** These authors contributed equally to this work (Omar Ramírez-Nuñez, Mariona Jové, Pascual Torres). To whom correspondence should be addressed at IRBLleida-UdL, Edifici Biomedicina I, Avda Rovira Roure 80 E25196 Lleida, Spain.

**Keywords:** Motor neuron, lipidomic, nuclear envelope, polyunsaturated fatty acids, subcellular lipidomics, phospholipase C ßI, protein kinase CßII, alkyldihydroxyacetonephosphate synthase

## Abstract

In this pilot study, we show that nuclei in spinal cord from ALS patients exhibit a differential lipidomic signature. Among the differential lipid species we could annotate 41 potential identities. These comprise membrane-bound lipids such as phosphatidylethanolamines–including plasmalogens- and phosphatidylcholines but also other lipid classes such as glycosphingolipids, diacylglycerols, and triacylglycerides (potentially present as nuclear lipid droplets). These results were orthogonally validated by showing loss of alkyldihydroxyacetonephosphate synthase (AGPS), a key peroxisomal enzyme in plasmalogen synthesis, both in ALS necropsy samples, in human motor neurons derived from iPSC from ALS patients and in hSOD-G93A transgenic mice. Further, diacylglycerol content changes were associated to ALS-linked variations in related-enzymes, such as phospholipase C ßI (PLCßI), the source of nuclear diacylglycerol, and protein kinase CßII (PKCßII), whose function partially depends on nuclei concentration of diacylglycerol. These results point out for not only a role of nuclear membrane lipids but also to lipids present in the nucleoplasm, suggesting an undisclosed role for this part of the subcellular lipidome in ALS pathophysiology.

## INTRODUCTION

Amyotrophic lateral sclerosis (ALS) is a devastating neurodegenerative disease, where patients exhibit a loss in motor neuron number. This cell demise has been associated to a number of cellular events common to other age-related neurodegenerative conditions. These include the accumulation of modified proteins, such as Tar-DNA binding of 43 kDa (TDP-43) and ERK, in non-physiological subcellular locations [1]. Interestingly, these proteins play essential roles in nucleus, so their presence out of nucleus might be a consequence of an impairment in normal nucleocytosolic traffic. Indeed, it is known that this physiological trait is affected in ALS[2]. Nucleocytosolic transport depends on a series of proteins such as Ran-GTPases as well as the nuclear pores function. Of note, these nuclear pores are embedded into the nuclear envelope lipid bilayers. Further, recent data demonstrate alterations in nuclear membrane in ALS models [3] Lipids are key players in cellular physiology. Lipid composition is an essential feature of neural tissue. Recent data from our group and others have revealed that in ALS an extensive array of changes involving tissue lipidome is present. Indeed, in experimental models of ALS, dietary changes in lipid composition induce changes in phenotype of this neurodegenerative condition. Furthermore, epidemiological data demonstrate that the dietary intake of selected lipids (such as those belonging to n-3 family) is a relevant factor in predicting the incidence of this condition[4].

To the date, the lipidome of nuclear envelopes has been focus of relative attention in some conditions [5]. However, whether it is affected in ALS is currently unknown. To shed light into this question, we designed a pilot study to analyze samples from spinal cord of ALS patients, and age and gender matched individuals, to explore if ALS is associated to changes in nuclear lipidome(s). This has been achieved after nuclei enrichment from frozen tissues after necropsy and subsequent lipid analyses by liquid chromatography coupled to time-of-flight mass spectrometry. The results reveal an important change not only in typical membrane lipids but also affecting signaling molecules. The suggested pathogenic pathways have been validated independently by evaluating the levels and distribution of key proteins related with differential lipids, both in human samples and in a transgenic mice model overexpressing an ALS-related gene.

## METHODS

### Human and mice neuronal tissues

All human samples were obtained from the Institute of Neuropathology and HUB-ICO-IDIBELL Brain Bank following the guidelines of the local ethics committees. Extensive pathological studies were done for ALS diagnosis as previously described[6]. Samples lumbar spinal cord were from 2 males and 2 females aged between 57 and 79 years affected with typical neurological and neuropathological characteristics of sporadic ALS. The post-mortem delay between death and tissue processing was between 3 and 16h. Age- and gender-matched controls with no clinical evidences of neurological disease and with a normal neuropathological study were processed in parallel (see Table 1).

**Table 1.** Characteristics of tissue donors.

| **Gender** | **Age at death** | **Diagnosis** | **Disease onset** | **Disease duration** | **Postmortem delay** |
| --- | --- | --- | --- | --- | --- |
| M | 64 | ALS | Bulbar | 2 years | 16h 30 m |
| M | 57 | ALS | Bulbar | 2 years | 4h |
| F | 75 | ALS | Bulbar | 4 years | 4h 5m |
| F | 79 | ALS | Bulbar | 20 months | 2h 10m |
| M | 74 | - | - | - | 10h 50m |
| F | 78 | - | - | - | 5h |
| M | 66 | - | - | - | 4h 20m |
| M | 86 | - | - | - | 18h 15m |

We also employed lumbar spinal cord samples from male mice (150 d, n=5) from the strain B6SJL-Tg (SOD1-G93A)1Gur/J (from now on G93A) purchased at The Jackson Laboratories (JAX catalogue stock number 002726, Bar Harbor, MN, USA) and maintained in the B6SJL background, by male founder crossing with B6SJLF1/J. As controls we employed non-transgenic littermates. Mice care and housing have been previously described [7]. Animals, after being fasted overnight, were anaesthetized with 2.5% isoflurane in air and finally sacrificed. Spinal cords were rapidly excised, frozen in liquid N_2_ and stored at −80°C until further analyses.

For western-blot, samples were homogenized on ice in a buffer containing 180 mM KCl, 5 mM MOPS, 2 mM EDTA, 1mM diethylenetriaminepentaacetic acid, and 1 µM of freshly prepared butylated hydroxyl toluene at pH 7.3 using a homogenizer device (T10 basic UltraTurraX, IKA, Staufen, Germany). Protein concentrations were measured by the Bradford method. For immunohistochemistry, animals after sacrifice were exanguinated by perfusion, firstly with saline solution (4°C) and thereafter with ice cold 0.4% paraformaldeyde (Sigma-Aldrich, Sant Louis, MO, USA) solution (freshly prepared with pH 7.4 phosphate buffer). All experimental procedures were approved by the Institutional Animal Care Committee of IRBLleida and were conformed to the Directive 2010/63/EU of the European Parliament.

### Human Induced pluripotent stem (hIPS)cell-derived motor neurons

hiPS were obtained from NINDS Human Genetics DNA and Cell line Repository at Coriell Institute (Camden, NJ, USA #ND35663–with a mutation in *FUS*-, #ND42765–with a C9ORF72 expansion-, and #ND41865 – control cell line-) and grown in standard human embryonic stem (ES) cells media supplemented with 6ng/ml basic Fibroblast Growth Factor (bFGF) on inactivated mouse embryonic fibroblast (MEFs) (Sigma) as feeder cells. Differentiation towards motor neurons was carried by a modification of Du et al protocol[8]. Briefly, undifferentiated hiPSC cultures were digested with Accutase solution (Sigma-Aldrich) for 5 min, seed for 40 min in MEFs-conditioned hES media plus 10µM Y27632 (Cayman Chemicals, Ann-Arbor, MI, USA) to remove as many fibroblasts as possible, then the cell suspension was transferred into Geltrex®(Thermo Scientific)-coated tissue culture dishes for 24-48hrs until stem colonies were visible. Media was then changed to neuroepithelial media (NEPM) for 6 days, and then replaced sequentially to motorneuron progenitor (MNPs) media 1st and 2nd, 6 days in each of them. At this point, progenitors were either expanded on MNPs 2nd or further differentiated into neurospheres and motor neurons using motor neuron induction media and motor neuron maturation media respectively. Importantly, for neurosphere formation non-tissue culture dishes without coating were used. Both neuroepithelial and MNPs were split into new Geltrex®-coated dishes when 70-100% confluent and seed at 50-75 000 cells/cm² and always using Y27632 10µM during the 1st 24 hrs. Media was changed every other day.

After 6 days in motor neuron induction media, neurospheres were collected in a15 ml centrifuge tube, spin down 3.5 min at 0.1rcf, supernatant discarded and spheres digested for 5-7 min in Accumax solution (Sigma). With the help of a pipette, cells clusters were disaggregated until an homegenous single cell suspension was obtained. Cells were then washed with DMEM/F12 (Thermo Scientific) and resuspended in motor neuron maturation media for seeding, media was supplemented with Y27632, Laminin (2.5 µg per 1ml of media, Thermo-Fisher) and 0.1µM Compound-E (Stem Cell Technologies, Grenoble, France) during the first 3 days after plating in Poly-L-Lysine (Sigma) coated dishes, media change was done every 72 hrs. Seeding density was maintained at 50-100 000 cells/cm². The different media composition appears in the Table 2.

**Table 2.** Media used for human motor neuron differentiation from iPSC.

| **Component\Media** | **NEPM** | **MNP media**<br>**1st** | **MNP med**<br>**2<sup>nd</sup>** | **MN**<br>**induction**<br>**med** | **MN**<br>**maturation**<br>**media** |
| --- | --- | --- | --- | --- | --- |
| **DM/F12** | 20 ml | 20 ml |  |  |  |
| **Neurobasal** | 20 ml | 20 ml |  |  |  |
| **Neurobasal +** |  |  | 40 ml | 40 ml | 40 ml |
| **B27+ supplement** | 800 µl | 800 µl | 800 µl | 800 µl | 800 µl |
| **Ascorbic acid** | 0.1 mM | 0.1 mM | 0.1 mM | 0.1 mM | 0.1 mM |
| **L-Glutamine solution**<br>**(Thermo Scientific, #**<br>**25030024)** | 1:200 | 1:200 | 1:200 | 1:200 | 1:200 |
| **Non essential amino**<br>**acid solution (Sigma,**<br>**M7145)** | 1:100 | 1:100 | 1:100 | - | - |
| **CHIR** | 3 µM | 1 µM | 3 µM | 3 µM |  |
| **SB43** | 2 µM | 2 µM | 2 µM | 2 µM |  |
| **DMH1** | 2 µM | 2 µM | 2 µM | 2 µM |  |
| **Retinoic acid** |  | 0.1 µM | 0.1 µM | 0.5 µM | 0.5 µM |
| **Purmorphamine** |  | 0.5 µM | 0.5 µM | 0.1 µM | 0.1 µM |
| **Valproic acid** |  |  | 0.5 mM |  |  |
| **Compound E** |  |  |  |  | 0.1 µM |
| **CNTF** |  |  |  |  | 20 ng/ml |
| **IGF1** |  |  |  |  | 20 ng/ml |
| **Penicillin/Streptomycin**<br>(Thermo Scientific #<br>15140122) | 1:400 | 1:400 | 1:400 | 1:400 | 1:400 |

### Nuclear enrichment

Nuclei were extracted from portions of the human spinal cord according to a variation of the Blobber and Potter protocol [9]. We started from about 0.270 g of frozen tissue, which was allowed to thaw at 4 °C. All subsequent steps, unless noted otherwise, were performed at this temperature. Once thawed, samples were cut into very small pieces with a scalpel and transferred to a 2 mL eppendorff, where the tissue was homogenized in two volumes (w/v) of 0.25 M sucrose in cold TKM (0.05 M Tris, 0.025 M KCl. MgCl2 0.005 M) and commercial protease and phosphatase inhibitors (78440, Thermo Scientific). The homogenization was first performed with a Polytron® mechanical homogenizer with a plastic plunger with approximately 20-40 strokes. The homogenate was then passed to a Dounce-type homogenizer with a glass tight plunger (20-40 strokes). At the end of this step, the homogenate (540 μL) was transferred to an ultracentrifuge tube (Beckman Coulter, 344057), and 1080 μL of 2.3 M sucrose in TKM were added, with the contents of the tube were carefully mixed by inversion. A pipette was then introduced to the bottom of the centrifuge tube and 540 μL of 2.3 M sucrose in TKM were carefully added, ensuring that well-defined phases of sucrose were formed, where the homogenate remained at the top. The ultracentrifuge tubes were centrifuged at 124000 x g at 4 ° C for 30 minutes in Optima L-100XP ultracentrifuge (Beckman Coulter, SW 55 Ti rotor). The resulting pellet was suspended in 50 μL of TKM with protease inhibitors and commercial phosphatases (78440, Thermo Fisher Scientific). The presence of nuclei was confirmed by fluorescence microscopy, after staining with DAPI at 1 μg/mL. For comparison of nuclear enrichment, mitochondrial enriched fractions were isolated as indicated in Mitochondria Isolation Kit for Tissue (Ab110168, Abcam).

### Lipidomics of isolated nuclei

The lipids of the isolated spinal cord nuclei extracted post mortem of ALS patients and controls were extracted using methyl-tert-butyl-ether with the help of a sonicator, after adding deuterated internal standards representing major lipid classes[10]. The analysis was performed using a liquid chromatograph (Agilent 1290 UPLC) coupled to a mass spectrometer (Agilent 6520 ESIQTOF-MS) that allows the characterization of lipids and metabolites based in the exact mass (<10 ppm), isotopic spectrum and in retention times of internal standards marked isotopically and representative of the major lipid classes. Lipid species were separated through reverse phase chromatography (Waters Acquity HSS T3 × 0.1 mm × 100 mm at 55 ° C), in suitable gradients and previously described [10]. The eluate was monitored in the Q-TOF system by electroespray in positive and negative ionization mode, collecting data in a range of m/z from 100 to 3000, with ionization source at 120°C and using N2 as a solvation gas at 250 °C (800 L/h). The exact mass adjustment is performed by joint infusion in reference spray of purine and HP9982, with exact masses with errors less than 0.5 ppm. Tandem mass spectrometry was performed as described[10].

After the extraction of the molecular characteristics, the analysis of the results was performed using the Mass Profiler software (Agilent, version for mass spectrometry). All samples were coded and analyzed in double blind mode. After the data collection, they were decoded and the relevant statistical analyses, both univariate and multivariate, by the R package OrbisMet (available in http://hdl.handle.net/10609/65305)

### Microscopy of isolated nuclei

The isolated nuclei were fixed in 4% paraformaldehyde (w/v) in PBS at a 1/3 (nuclei solution/fixation solution) at room temperature for 15 minutes. Subsequently, 20 μL of the mixture were placed on a round coverslip (12 mm), which was subsequently centrifuged (100xG, 3 min). At the end of the time, coverslips were taken with fine forceps and placed on a Parafilm®-covered support. The nuclei were incubated in a permeabilization and blocking solution (Triton X-100 0.1%, normal goat serum 10% in PBS) for 30 minutes at room temperature. Then, three gentle washes were carefully made with PBS. Coverslips were incubated at 4°C overnight (in a humid chamber) with the appropriate primary antibodies at the indicated final concentrations, dissolved in 1/10 (v/v) of blocking solution in PBS. After this incubation period, three washes were made with PBS and the samples were incubated with the appropriate secondary antibody dissolved in PBS and 1 μg / mL DAPI for nuclear staining, at 4 °C for 1 hour. Finally, 3 washes of 5 minutes each were made with PBS and the nuclei were mounted on slides with Fluoromount-G solution (00-4959-52, Thermo Fisher Scientific) and allowed to dry at room temperature. Nile Red at 10µg/mL in PBS was used to evaluate the presence of lipid droplets, by incubating coverslips containing fixed nuclei at 37°C for 30 min.

### Western Blot analysis

In case of human samples, protein from different fractions (total lysates, mitochondrial and nuclei enriched fractions) was extracted using radioimmunoprecipitation buffer with Halt Protease Inhibitor Cocktail (1X) (1861278, Thermo Fisher Scientific). After sonication, protein quantification was performed with Bradford assay. Both for human and mice samples, 15 µg of protein were loaded onto a 12% acrylamide SDS-PAGE gel. Membranes were blocked with I-Block (T2015, Thermo Fisher Scientific) for 1 hour and incubated overnight with the antibodies and conditions listed in Table 2. After primary antibody incubation, membranes were washed 3 times with TBS-T 0.05% and incubated with secondary antibody for 1h. Immobilon™ Western Chemiluminiscent HRP Substrate (Merck Millipore, WBKLS0500, Billerica, MA, USA) was used for immunodetection. Membranes were stained with Coomassie Brilliant Blue G (27815, Sigma-Aldrich) for normalization. Specific bands were quantified with ImageLab v5.2.1 (Bio-Rad). Omission of primary antibody resulted in complete loss of signal.

### Immunofluorescence

Immunofluorescence was performed essentially as previously described [1]. Briefly, after tissue collection (lumbar spinal cord) in mice, samples were immersed for 24h in paraformaldehyde solution to assure preservation. Samples were thereafter immersed in 30% sucrose, (made in pH 7.4 phosphate buffer) for 48h to achieve cryopreservation. Tissue was encased in a cubic recipient (Peel-A-Way Disposable Embeding Molds-S-22, Polysciences Inc., Warrington, PA, USA), labelled and embedded in tissue freezing medium (Triangle Biomedical Sciences Inc., Newcastle, UK) for easy cutting and better preservation and finally frozen (−80°C). Sixteen µm wide sections of lumbar spinal cord were cut and carefully seeded along gelatin-coated slide. Selected sections were permeabilized with 0.1% Triton X-100 in PBS for 30 min at room temperature. Blocking of sections was performed with 5% normal goat serum in 0.1% Triton X-100 PBS at room temperature for 1h. In the case of human MNs, cells (grown in glass coverslips) were washed with PBS and then fixed with 3.7% paraformaldehyde for 10 min at room temperature. Cells were rinsed with PBS, permeabilised with 0.1% Triton X-100 in PBS for 30 min and subsequently blocked with 5% fetal horse serum at room temperature for 2 hours.

For immunofluorescence, slides (mouse spinal cord sections) or coverslips (human MNs) were incubated at 4°C overnight with the primary antibodies indicated on table 3. Next day, slides and coverslips were washed with PBS three times for 5 min at room temperature. Then, they were incubated at room temperature for 1h in darkness with corresponding secondary antibodies. Sections and cells were finally counterstained with 4,6-diamidino-2-phenylindole dihydrochloride (DAPI, 1 ug/ml) in PBS at room temperature for 10 min and, after three washes with PBS, mounted on slides with Vectashield (Vector Laboratories, Burlingame, CA, USA). In selected sections and coverslips, primary antibody was omitted to assure labelling specificity. Mounted slices were examined under a Fluoview 500 Olympus confocal laser scanning microscope (Olympus, Hamburg, Germany). For quantification tissue immunoreactivity images were analyzed with the Image J software (http://rsbweb.nih.gov/ij/) and human MNs images were evaluated using an ad-hoc designed Cell Profiler pipeline (http://cellprofiler.org), available in supplemental materials.

**Table 3.** Antibodies and conditions used.

| Target | Dilution | Reference |
| --- | --- | --- |
| PARP | 1:500 in TBS-T 0.05%<sup>1</sup> | sc-74470 (Santa Cruz Biotechnology) |
| Histone H3 | 1:500 in TBS-T 0.05% | SAB4500352 (Sigma-Aldrich) |
| Mitofusin 2 | 1:1000 in TBS-T 0.05% | M6319 (Sigma-Aldrich) |
| Porin | 1:1000 in TBS-T 0.05% | ab15895 (Abcam) |
| Lactate dehydrogenase | 1:1000 in TBS-T 0.05% | ab135396 (Abcam) |
| Nuclear Pore Complex Proteins<br>[Mab414] | 1:250 in PBS | ab50008 (Abcam) |
| Tubulin βIII | 1:250 in PBS | ab18207 (Abcam) |
| AGPS | 1:250 in TBS-T 0.05% for<br>western blot; 1:150 in PBS<br>for immunofluorescence | sc-374201 (Santa Cruz Biotechnology) |
| PLC βI | 1:250 in TBS-T 0.05% | sc-5291 (Santa Cruz Biotechnology) |
| PKC βII | 1:250 in TBS-T 0.05% | sc-13149 (Santa Cruz Biotechnology) |
| SCP-2 | 1:1000 in TBS-T 0.05% | hpa027135 (Atlas Antibodies) |
| PMP70 | 1:250 in PBS | Ab3421 (Abcam) |
| Mouse IgG, Alexa Fluor® 488<br>conjugate | 1:800 in PBS | A-11001 (Thermo Fisher Scientific) |
| Rabbit IgG, Alexa Fluor® 546 conjugate | 1:800 in PBS | A-11010 (Thermo Fisher Scientific) |
| Rabbit IgG, horseradish peroxidase conjugate | 1:50000 in TBS-T 0.05% | 31460 (Thermo Fisher Scientific) |
| Rabbit IgG, horseradish peroxidase conjugate | 1:30000 in TBS-T 0.05% | NA931V (Sigma) |

<sup>1</sup> TBS-T 0.05%: Tris Buffered Saline with Tween-20 0.05%

**Table 4.** Differential lipids in nuclei isolated from ALS patients in comparison to non-ALS.

| Potential ID<sup>1</sup> | P value<sup>2</sup> | Change(ALS vs non-ALS) |
| --- | --- | --- |
| Diacylglycerols |  |  |
| DG(34:2)<sup>3</sup>[iso2] | 0,017499 | down |
| DG(46:3)/CE(24:0) | 0,027414 | down |
| DG(36:0)<sup>3</sup> | 0,023379 | down |
| DG(36:2)<sup>3</sup> | 0,010411 | down |
| DG(39:7)[<sup>3</sup>/PA(O-32:0) | 0,007198 | up |
| DG(35:1)<sup>3</sup> | 0,025619 | up |
| DG(36:0)<sup>3</sup> | 0,001003 | up |
| DG(38:0)<sup>3</sup> | 0,002498 | up |
| DG(39:6)<sup>3</sup> | 0,034444 | up |
| Glycerophospholipids |  |  |
| PA(18:0) | 0,024149 | down |
| PA(O-36:3)/PG(O-29:0) | 0,043124 | down |
| PG(P-40:2)/PG(O-40:3) | 0,027358 | down |
| PS(O-40:1)/PS(P-40:0)/ PS(39:1)/ | 0,015925 | down |
| PE(P-42:1)/ PE(O:42:2)/PE(42:1)/GlcCer(d36:0)/PC(39:1) | 0,029046 | down |
| PA(O-32:0) | 0,024342 | up |
| PA(P-37:0)/PG(32:0)/PA(38:5) | 0,018452 | up |
| PA(P-35:1) | 0,018294 | up |
| PC(O-24:0) | 0,0235 | up |
| PE(42:10)/PI(31:1) | 5,35E-05 | up |
| PE(O-37:1)/ )/ PC(P-33:2)/PE(P-36:2)/PE(O-36:3)/Cerebroside B/<br>GlcCer(35:2)/CerP(42:2) | 0,024172 | up |
| PE(P-38:3)/PE(38:3)/PC(O-35:4)/PE-NMe2(36:2)/PE(O-38:4 | 4,76E-04 | up |
| PG(33:2) | 0,025597 | up |
| PG(O-38:1)/PG(P-38:0)/PG(38:0)/TG(48/8) | 0,025968 | up |
| PG(P-37:1)/PG(O-37:2) | 0,022001 | up |
| PS(P-38:4)/PS(O-38:5)/PI(O-32:0)/PC(37:4)/PE(40:4) | 0,01172 | up |
| PS(P-38:4)/PS(O-38:5) | 0,025977 | up |
| PE(P-38:3)/PE(38:3)/PC(O-35:4)/ PE(O-38:4)/Cerebroside C/ | 0,011598 | up |
| Triacylglycerols |  |  |
| TG(46:1) | 0,029092 | down |
| TG(48:2) | 0,028442 | down |
| TG(50:3)/PG(41:0) | 0,027652 | down |
| TG(60:7) | 0,019288 | down |
| TG(43:3)/PE-Cer(39:1)/SM(36:1)/PA(P-39:0)/PA(O-39:1) | 0,027467 | down |
| TG(54:3) | 0,01669 | up |
| TG(65:12) | 0,001038 | up |
| TG(50:0) | 0,027069 | up |
| TG(59:9) | 0,00401 | up |
| TG(65:2) | 0,001892 | up |
| Sphingolipids |  |  |
| Cer(d35:0) | 0,027563 | down |
| GlcCer(d42:1)/PC(38:1)/PE(41:1) | 0,026204 | down |
| Cer(46:0) | 0,007346 | up |
| LacCer(d34:0)/PC(42:5)/Ins-1-P-Cer(40:1)/C22 Sulfatide | 0,025398 | up |

<sup>1</sup> LIPID-MAPS nomenclature, identification has been proposed on basis of isotope distribution and identity of retention times with those of representative standards, except in <sup>3</sup> where identity was confirmed by MS/MS; <sup>2</sup> p value after T test for differences

### Statistical analyses

All statistics were performed using the Prism software (GraphPad Software, San Diego, CA, USA). Differences between groups were analyzed by the Student’s t tests or ANOVA after normality of variable distribution was ensured by Kolmogorov-Smirnov test. The 0.05 level was selected as the point of minimal statistical significance in every comparison

## RESULTS

The described method (Figure 1a) leads to a fraction enriched in nuclei, derived from frozen tissues. The nuclei present in this fraction exhibit a preserved nuclear morphology (DAPI positivity with high nucleoporin content–Figure 1b). Further, a high content of nuclear specific proteins was demonstrated by western blot analyses (Figure 1C). The potential presence of membrane containing organelles, such as mitochondria, was excluded, based on the content of mitochondrial membrane markers such as porin or mitofusin (Figure 1c).

**Fig. 1.**
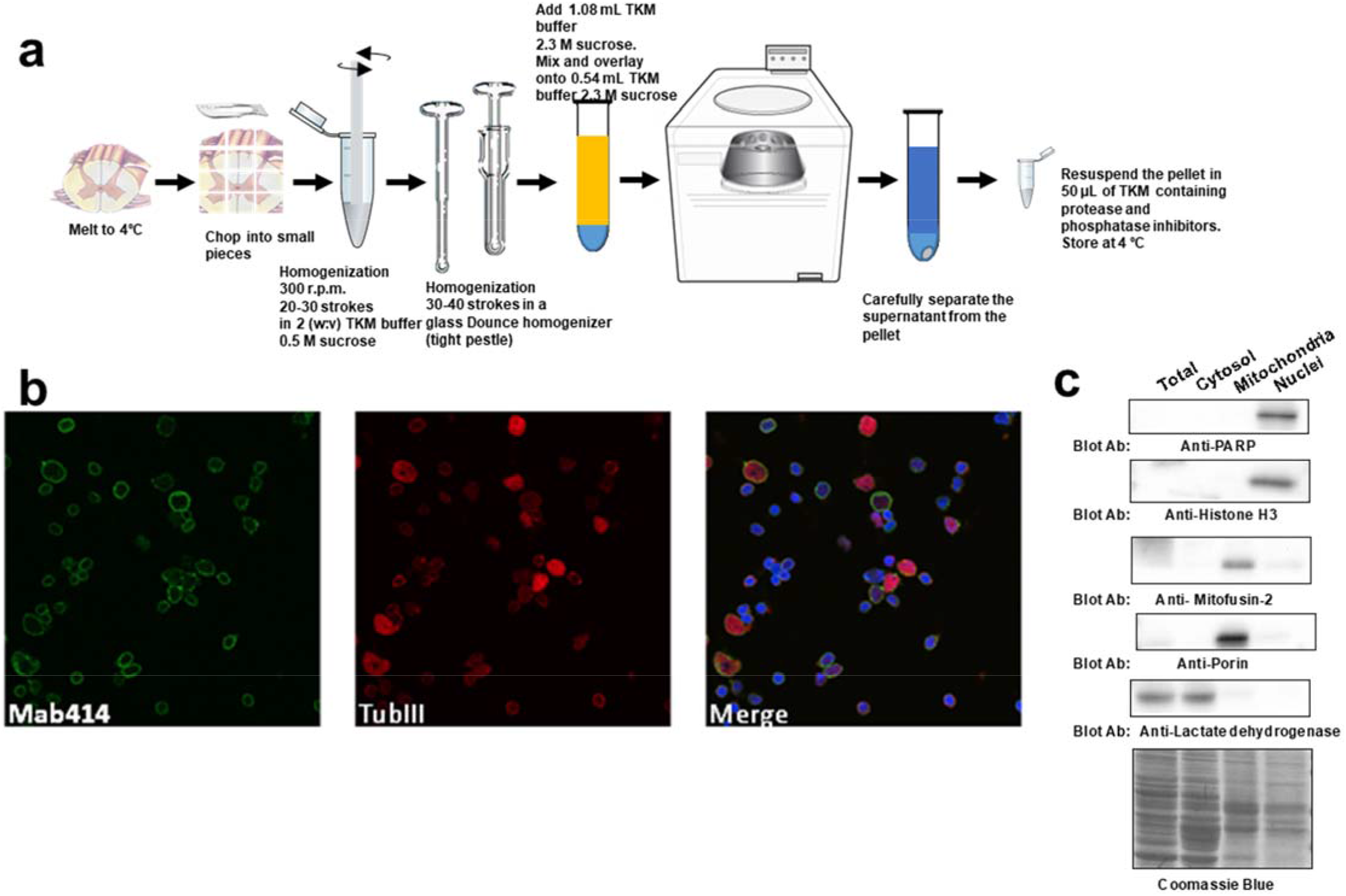
Whole nuclei could be enriched from frozen spinal cords. a) Schematic procedure for obtention of a nuclei enriched fraction departing from frozen human spinal cords. b) Immunocytochemical evidence for nuclei integrity and heterogeneity, as shown by immunolabeling of nucleoporin (Mab414), by presence of neuron specific tubulin ß III (TubIII) and DAPI possitivity (in blue). c) Nuclei enriched fraction is depleted from cytosolic and other membrane containing organelle (such as mitochondria), as indicated by lack of immunoreactivity for mitochondrial markers (porin and mitofusin) and selective improvement in nuclear antigens (poly-ADP-ribose polimerase (PARP) and histone-H3)

Lipidomic analyses disclosed significant differences between nuclei from ALS and age-matched healthy controls. Thus, principal component analyses (PCA) explained more than 50% of total variances (either in negative or in positively ionized lipids) of present samples in a model of three components (Figure 2a). Further, partial-least square models offered an accuracy in predicting the groups *ca* 100%, with more than 60% of total variance in lipidomic profiles being explained in three major components, demonstrating the robustness of the model (Figure 2b). As shown by hierarchical clustering analyses (Figure 2c), marked differences were present among the lipidomic profiles of the resulting nuclei enriched fractions.

**Fig. 2.**
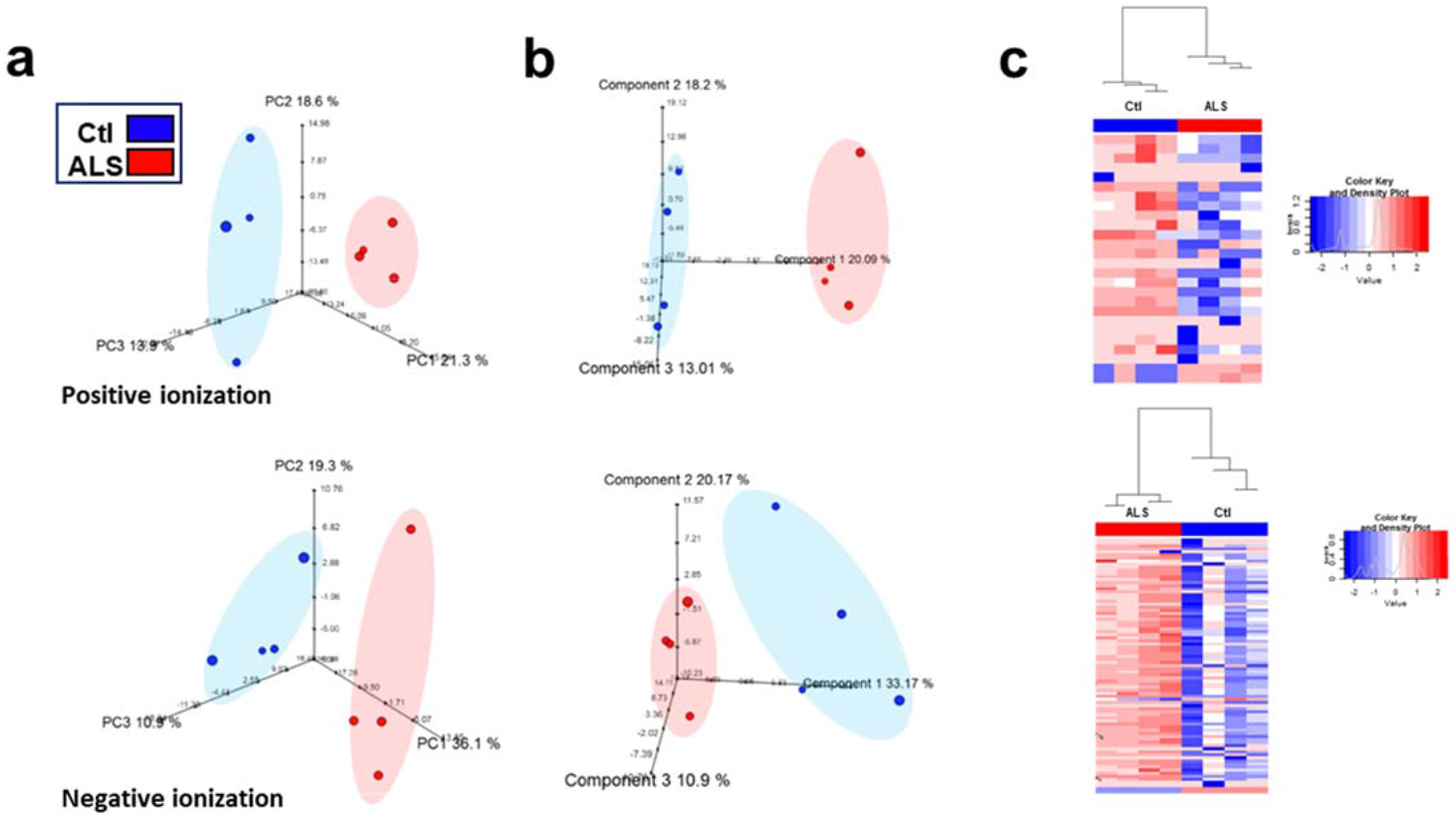
Nuclei from ALS patients exhibit a specific lipidome. Multivariate analyses including PCA (a), partial least square-discriminant analyses (b) and hierarchical clustering analyses (c, using lipids significantly affected by ALS status) demonstrate that nuclei from ALS patients exhibit a specific lipidomic profile. Upper panels are obtained by using lipids ionized in positive ionization, while as lower panels show the results using negative ionization

Univariate analyses (T test) demonstrated the presence of 151 differential molecules (99 in positive ionization and 52 in negative ionization, Supplemental table 1, p values between 9.46×10E-15 and 0.05). Among these, putative identities, based on isotope distribution pattern and retention time identity with internal standards representative of lipid major families, were found for 41 different molecules (25 upregulated in ALS samples; Table 3), with p values ranging from 5.35×10E-5 to 0.05. Potential identities (comprising isobaric molecules) included 8 ceramides, 9 diacilglycerol, 8 phosphatidylcholines,15 phosphatidylethanolamines, 9 phosphatidic acid, 7 phosphatidylserines, 2 phosphoinositides and 11 triacylglycerides.

After tandem mass spectrometry, we were able to confirm the identity of diacylglycerol molecules (DAG). One of the mediators of DAG is PKCßII. In line with altered levels of DAG, decreased levels of PKCßII were measured in both spinal cord lysates from ALS patients (Figure 3a) and lysates from G93A mice (Figure 3b), in comparison to adequate controls. Further, as DAG can be the result of the activation of an specific PLC isoform (ßI), we measured its levels in the same samples. Though immunoreactivity was near to marginal in human spinal cord lysates, clear decreases could be also measured in G93A spinal cord lysates (Figure 3c and 3d).

**Fig. 3.**
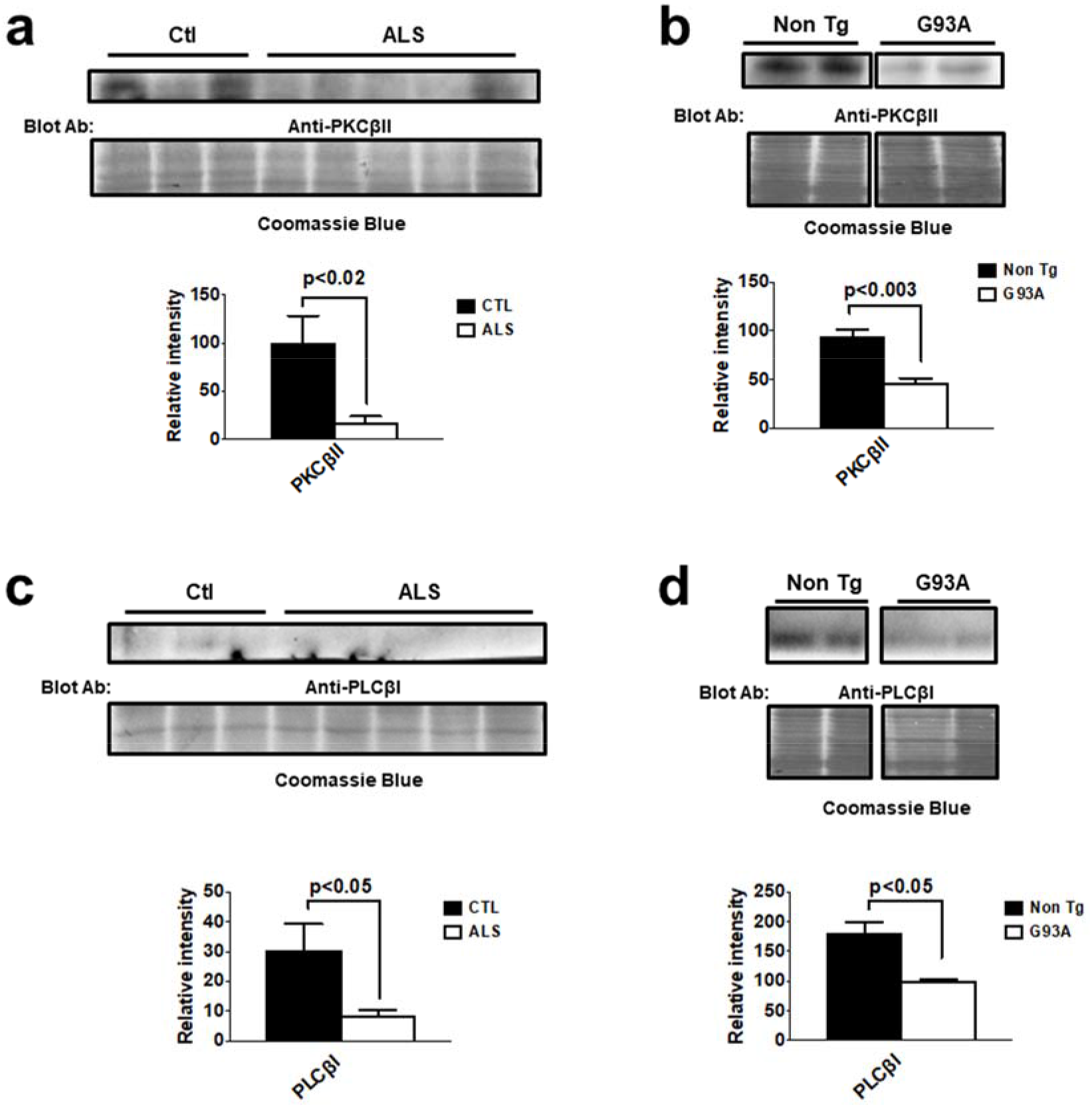
Changes in nuclear DAG concentrations suggested by lipidomics are associated with alterations in PLCßI and PKCßII levels in ALS. Representative western-blot analyses of PKCßII in spinal cord lysates from ALS patients (a) and G93A mice (at terminal stage, 150 days, b) show that protein concentration is decreased in comparison with adequate control. Similarly, western-blot analyses of PLCßI are also decreased in spinal cord lysates from ALS patients (c) and G93A mice (d). In all panels, upper part shows representative western-blot images with lower part demonstrating quantification by densitometry, after normalization by Coomassie blue staining. Differences between groups were analyzed by Student’s t test.

Similarly, as differential lipids also comprised several plasmalogens, we explored the potential changes of one of the rate-limiting enzymes for plasmalogen synthesis, AGPS [11], a peroxisomal enzyme. The results indicate that ALS could influence AGPS levels in spinal cord. Thus, western-blot analyses (Figure 4a) demonstrate significantly decreased levels in spinal cords of G93A mice in comparison with non-Tg animals. In contrast, in lysates from human samples, we were not able to reproduce this fact (Figure 4b). Noteworthy, in human motor neurons derived from IPs, the anti-AGPS immunroeactivity in two cell lines from known ALS-causing mutations (*FUS* and C9orf72 expansions) was lower than that from control cell line (Figure 4c). Further, there was a significant correlation between nuclear perimeter and AGPS levels in both control and C9orf72 cell lines, but not in *FUS* mutants. Confirming these data, in mice spinal cord, this protein was preferentially present in motor neurons, whose levels were severely decreased in 150 days old G93A mice (Figure 4d).

**Fig. 4.**
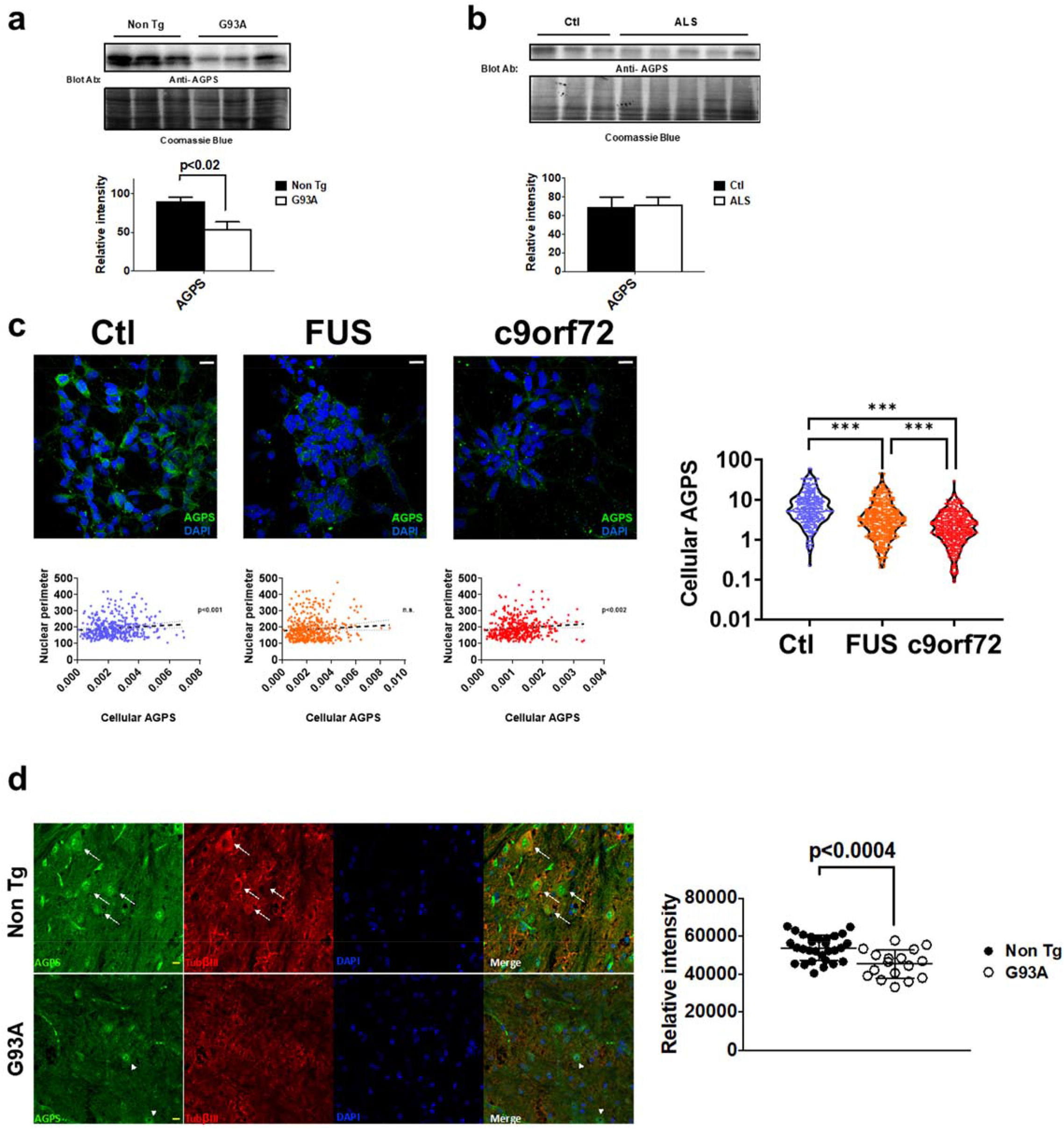
Changes in nuclear plasmalogen concentrations suggested by lipidomics are associated with alterations in AGPS in human motor neurons and in G93A mice. Representative western-blot analyses of AGPS in spinal cord lysates from G93A mice (at terminal stage, 150 days, a) and ALS patients (b) show that protein concentration is decreased in G93A mice in comparison with adequate controls. c) confocal imaging of hIPs-derived motor neurons demonstrates decreased levels in cells derived from *FUS* mutants and C9orf72 IPs, as shown by quantiatiative analyses (see right panel in c); further, there was a significant correlation between AGPS cell content and nuclear perimeter (see below panels from each picture) both in control and in C9orf72 cells, but this was lost in *FUS* mutated cells. d) shows that confocal immunohistochemistry analyses of ventral regions of lumbar spinal cord section demonstrate a strong reactivity of AGPS in motor neurons in non-transgenic mice (arrows) in comparison with G93A mice (arrowheads). Motor neurons were identified by size, nuclear morphology and tubulin ßIII immunoreactivity. Right panel in d shows quantitative analyses of AGPS immunoreactivity in these cells, compatible with decreased expression of this protein. In a) and b) panels, upper part shows representative western-blot images with lower part demonstrating quantification by densitometry, after normalization by Coomassie blue staining. Differences between groups were analyzed by Student’s t test (a and d) and by ANOVA followed by Bonferroni post-hoc analyses in c. Bars in c and d are 10 µM long.

Furthermore, some of the potential identities of the differential lipids are alkyl-phospholipids, whose synthesis is also dependent on peroxisome function. To evaluate whether changes in AGPS and in alkyl-phospholipids are due to a potentially specific peroxisomal defect, we evaluated PMP70 distribution in G93A mice. Results (Figure 5a) show that despite PMP70 enriched vesicles were present in many remaining motor neurons in G93A mice spinal cord, no general increases were evident by western-blot. Since we also detected differential phospholipid compositions in nuclear isolates, we evaluated SCP-2 expression. SCP-2 is a peroxisomal/ER resident protein implicated in phospholipid synthesis[12], particularly phosphatydilethanolamine and phosphatidylserine. In line with differential phospholipids found in nuclei isolated from ALS samples, we found a marked rearrangement of SCP-2 immunoreactivity in G93A mice. As shown in figure 5c, in control mice, SCP-2 expression was preferentially found around the nuclei, this was not the case in many motor neurons from G93A mice, with redistribution on this immunoreactivity towards vesicles with no perinuclear enrichment. Further, levels of SCP-2 were significantly increased, as shown by western-blot analyses.

**Fig. 5.**
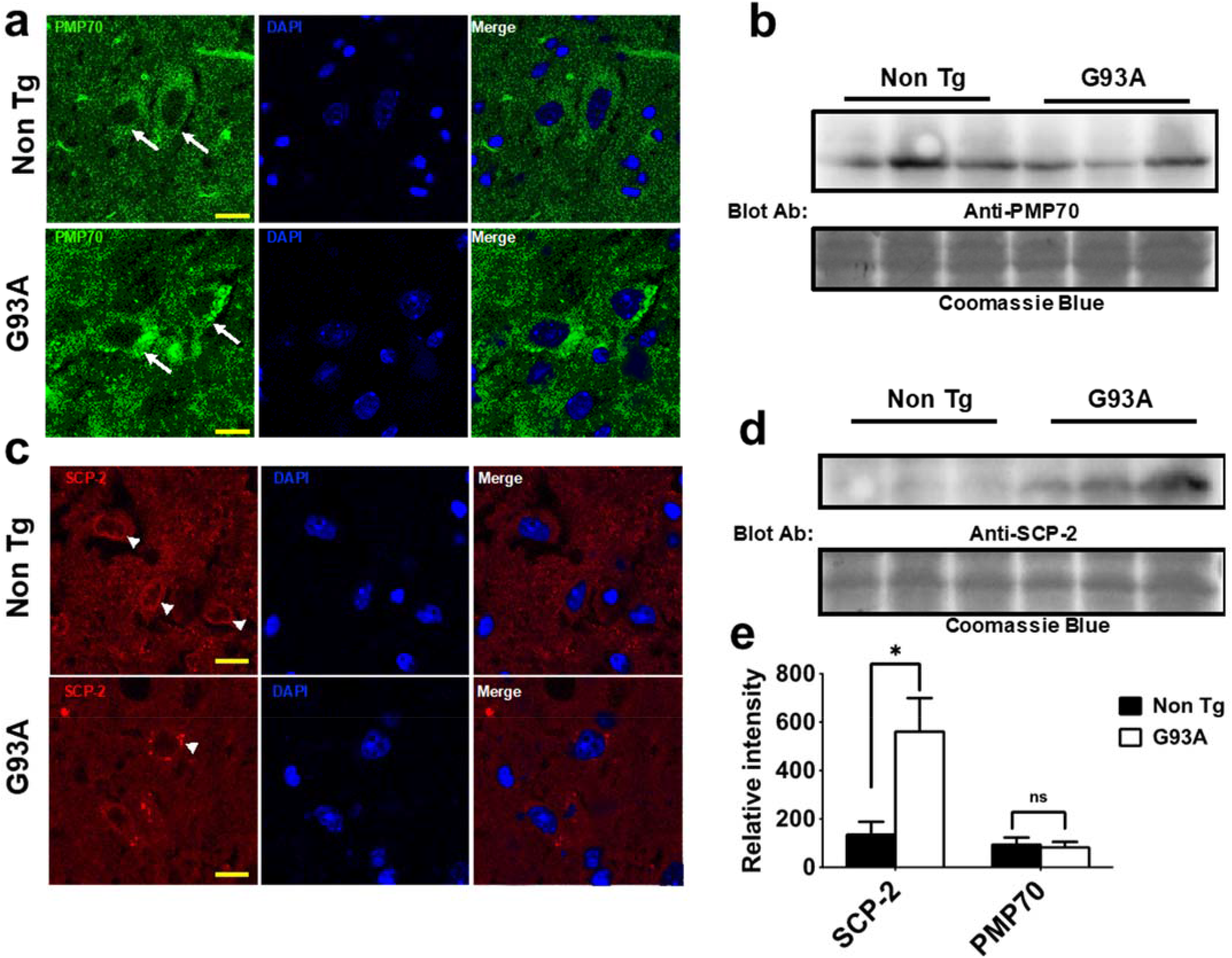
Changes in nuclear phospholipid and ether-phospholipids concentrations suggested by lipidomics are associated with alterations in SCP-2 in G93A mice. a) Confocal immunohistochemistry analyses of PMP70 in ventral regions of lumbar spinal cord sections, with an enrichment in PMP70 in motor neurons in G93A mice (at terminal stage, 150 days, arrows) in comparison with non-transgenic mice. Motor neurons were identified by size and nuclear morphology. b) As evidenced by western-blot analyses of PMP70 in spinal cord lysates, G93A mice do not show a general increase in PMP70 (quantification in e, after normalization by Coomassie blue staining). In line with potential ER/peroxisomal changes, immunoreactivity of SCP-2 is severely affected by G93A overexpression (c). While in non-Tg specimens, SCP-2 formed a perinuclear rim, this pattern is lost in G93A with enhancement in vesicular structures, specifically found in motor neurons. This is accompanied by an increase in the total lysate concentration of SCP-2, as shown by western blot analyses (in d and e). Differences between non-Tg and G93A were analyzed by Student’s t test. Bars in A and C are 10 µM long.

Finally, across the differential lipids in enriched nuclear fractions we detected molecules compatible with triacylglycerides. As these are usually found in lipid droplets[13], we explored if lipid droplets could be found along the purified nuclei. As shown in figure 6, Nile Red stained vesicles could be found associated with the nuclei and even inside.

**Fig. 6.**
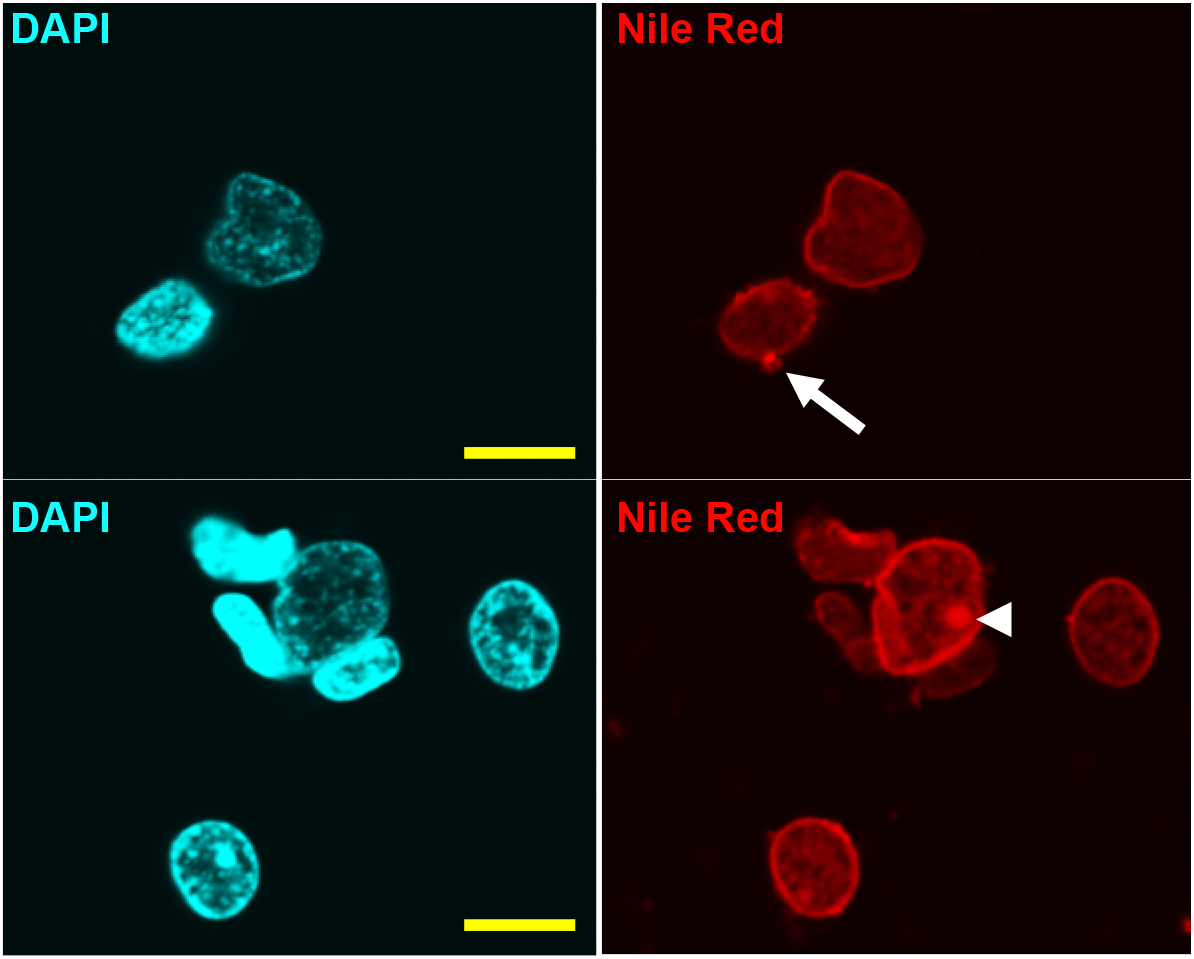
Isolated nuclei show structures compatible with lipid droplets. Confocal image of isolated nuclei, showing Nile Red fluorescent vesicles both in the periphery of the nuclei (upper panel) or even inside the nuclear matrix (lower panel). Bars are 10 µM long.

## DISCUSSION

The analysis of the lipidomic profile of the isolated nuclei of post tissue mortem of ALS patients and controls demonstrate a differential profile between control group and the pathological group. To the best of our knowledge, this is the first report indicating a difference in this subcellular parameter in this devastating neurodegenerative disease. There is not a large number of references on cellular membrane alterations in ALS and their models, although in previous studies of our group [6, 7], demonstrate severe changes in tissue lipidomes in ALS and ALS models, depending on disease stage and anatomical location.On the other hand, it should be remembered that one of the forms of familial ALS (subtype 8), is due to a mutation in the VAP-B protein, involved in intracellular membrane trafficking [14]. Thus, it can be expected that the composition of the intracellular membranes is a relevant factor in pathophysiology of ALS.

Specifically about the nuclear envelope, it has been recently described that failures in nucleocytosolic transport, depending in part from importins, are present in ALS [15]. The interaction between the lipids of the nuclear membranes and the nucleoporins–integral components of nuclear pore-as well as their contribution to cellular homeostasis is a field largely unknown. To perform their function, nucleoporins need to be anchored adequately at the fusion points between the nuclear membranes of external and internal nuclear envelope[16]. There are highly specialized regions of specific nucleoporins (such as Pom121, GP201) structurally dependent on phospholipids where these proteins are anchored into [17, 18]. In addition to these transmembrane proteins, other proteins, such as members of the NUP107-160 complex contain domains that interact with the nuclear membranes[19, 20]. Our results are compatible with changes in the composition of the membranes that make up the nuclear envelope, since phospholipids involved include phosphatidylethanolamine, phosphatidylcholine and phosphatidylinositol, all membrane components. Noteworthy, it shall be reminded that nuclear envelop leaflets are in continuity with endoplasmic reticulum membranes. Therefore, changes in nuclear envelope can be related with dysfunctions in this membranous organelle, already characterized in ALS[3,6,21]

DAG in nucleus has a signaling role, reviewed in [21], and it is derived from the activity of a PLCß1 isoform. In line with altered homeostasis of nuclear DAG in ALS, we detected different amounts of PLCß1 in tissue lysates from G93A mice and potentially from ALS patients. This enzyme is present in neuronal cells in rat brain [22], partially present in nuclei. Further, it is known that this enzyme is sensitive to endoplasmic reticulum stress[23], a well described pathogenic mediator in ALS[6]. Interestingly, a loss in PLCß1 has been also described in other neurodegenerative conditions linked to protein aggregation, such as Pick’s disease [24] and Creutzfeld-Jacobs disease[25]. Furthermore, PLCß1 loss induces an age-dependent loss of neurites and neurodegeneration[26]. Nonetheless, it is known that metabotropic glutamate receptors regulate synaptic transmission through the stimulation of PLCß1 [27]. Of note, activation of metabotropic signaling has been proposed as novel potentially therapeutic strategy in ALS[28]. DAG build-up in nucleus leads to PKCßII nuclear translocation[21]. For this reason, we evaluated whether, in line with altered DAG and PLCß1 levels in ALS, this disease is associated with changes in PKCßII. The results indicate that, both in ALS patients and in G93A mice, PKCßII levels are diminished. Up to date, no previous evidence of its involvement in this disease were known, except with the fact that PKC levels were increased in ALS patients [29]. The involvement of the potential loss of PKCßII in ALS pathogenesis can be diverse, as this protein interacts with mitochondria[30], and with autophagic function[31], to name a few, and both elements are involved in ALS [32, 33]

Other changes in lipid components of nuclei include plasmalogens. These membrane glycerophospholipids contain a fatty alcohol with a vinyl-ether bond at the sn-1 position, and they are enriched in polyunsaturated fatty acids at the sn-2 position of the glycerol backbone. For this reason, they have been involved in the redox homeostasis of membranes[34], and its Previous data have characterized plasmalogens in the nuclear envelope of postmytotic tissues such as myocardium [35]. Peroxisomes contribute to plasmalogen synthesis, and previous data of our group describe alterations in peroxisomal enzymes in ALS tissues [36]. Whether the loss of plasmalogen present in ALS nuclei is potentially derived from this peroxisomal disturbance and whether it contributes to ALS pathophysiology is still unknown. Nonetheless, AGPS intensity was strongly diminished in G93A mice motor neurons, supporting a potential involvement of this pathway in the pathophysiology of the spinal cord dysfunction present in these mice.

Among differential lipids, other classes are also present. These include triacylglycerides, which could be explained by changes in the so-called lipid droplets, recently described in the nucleoplasm[37] where it may derive from inner nuclear membrane[38]. Since lipid droplet physiological role in nuclei is mainly unknown, we cannot infer its pathophysiological significance in our samples. Nonetheless, recent data disclose that glial cells accumulate lipid droplets as a response to excitotoxicity[39]. However, they may also play a role in neurodegeneration in lipid peroxidation-prone environments[40]. Our finding that no general effect (not all triaclyglycerides increase or decrease) is present, may be derived from potentially heterogeneous cellular source of the nuclei.

Similarly, another group of lipids with potential relevance are sphingolipids. Previous findings in the field suggest the importance of these lipids in ALS. In the G93A model, alterations are demonstrated in tissue levels during the development of the disease. Further, the administration of GM3, one of its members, leads to an improvement in the clinical course of the disease [41]. In a close relationship, it has been also shown that the transcription of glucosylceramide synthase is necessary for optimal protection against denervation, and that its inhibition worsens the clinical course in another model of the family ALS, the mouse hSODG86R [42]. Our results are fully compatible with these data, as we found significantly decreased levels of two molecules compatible with glucosylceramide identity. As these molecules are functionally involved in cell signaling, autophagy and determination of lipid membrane functional domains [43, 44], its potential role in ALS pathophysiology merits further investigation.

We acknowledge, as a limitation of our work, the low number of individuals used. However, even with such a limitation, the profiling of a high number of nuclei is possible, and allows to distinction of clear lipidomic profiles. Further, the high heterogeneity of nuclear structures does not permit to attribute these changes to a specific cell subtype. Similarly, we do not know if freezing procedures have led to a selective enrichment of given cellular types or nuclei. Thus, we cannot exclude that specific nuclei can be more resistant to postmortem changes and criopreservation. However, we assume the fact that nuclei enrichment is robust, based on the absence of mitochondrial markers–major contaminants in similar procedures-, and the morphological and biochemical evidences for nuclear presence. Despite these limitations, changes in two lipid families (plasmalogen and DAG) were independently validated by exploring their potential molecular basis. We think that the combination of the described technologies (lipidomic analyses applied to subcellular fractionation of pathologically relevant samples) could pave the way for the discovery of novel biomarkers and pathophysiologically relevant pathways.

## DECLARATIONS

### Ethics approval and consent to participate

Samples from human individuals were obtained from the Institute of Neuropathology and HUB-ICO-IDIBELL Brain Bank following the guidelines of the local ethics committees. Experiments with mice were approved Institutional Animal Care Committee of IRBLleida and were conformed to the Directive 2010/63/EU of the European Parliament.

### Consent for publication

Not aplicable

### Availability of data and material

The Orbismet, the R package employed for analyses of lipidomic data during the current study are available in the UOC repository, at in http://hdl.handle.net/10609/65305

### Authors’ contributions

OR-N,MJ and PT optimized the nuclei isolation, and performed western-blot analyses. MJ and JS performed lipidomic data, designed R packages and analyzed these data. RR-G,PA-B, LF and CR generated transgenic mice, maintained iPSc and characterized motor neurons and derived results. VA,MP and JB performed general data analyses. RP, IF and MP-O planned the experiments, curated data, drafted the manuscript and prepared figures. All authors read and approved the final manuscript.

## Acknowledgements

We are indebted to tissue donors and their families.

---

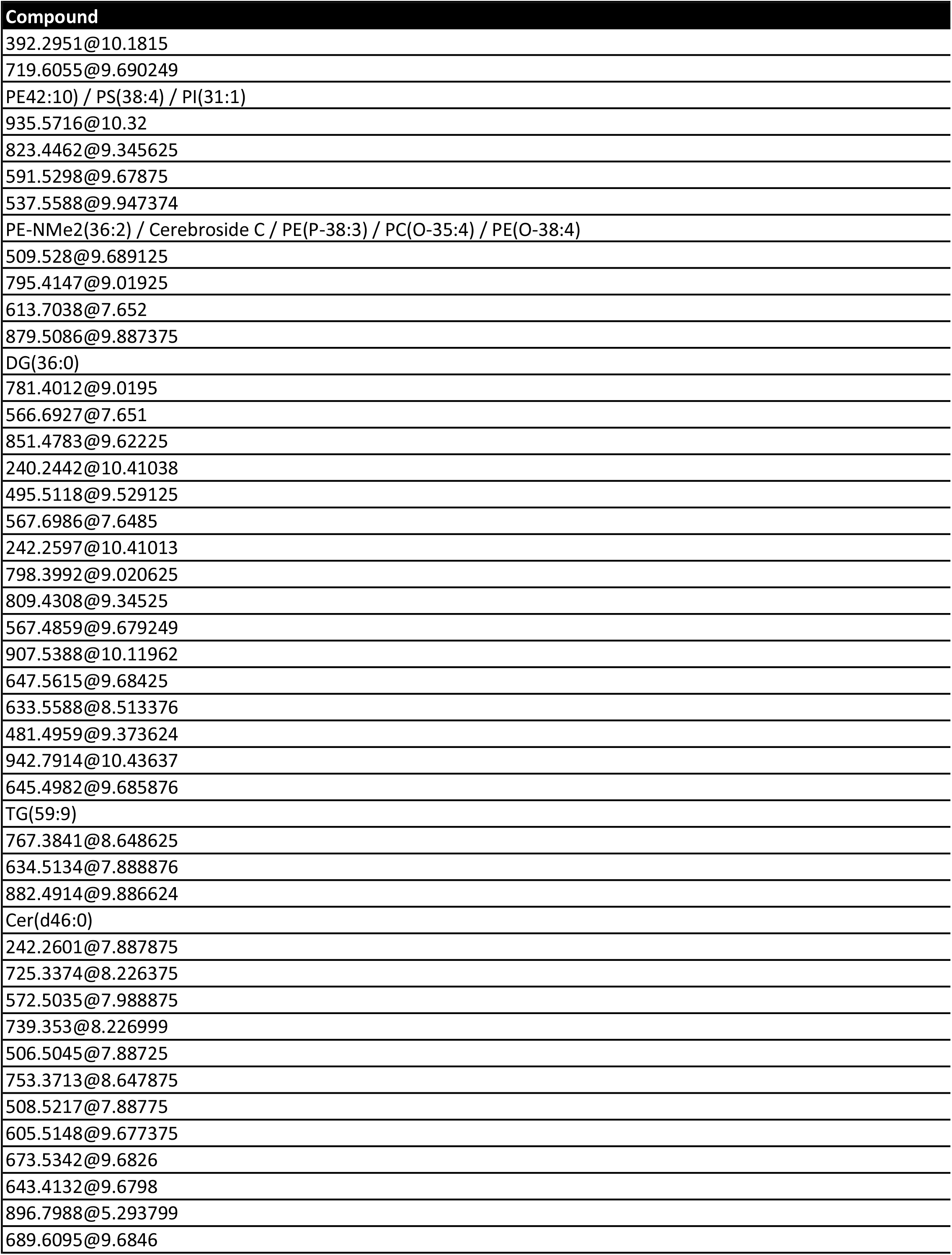
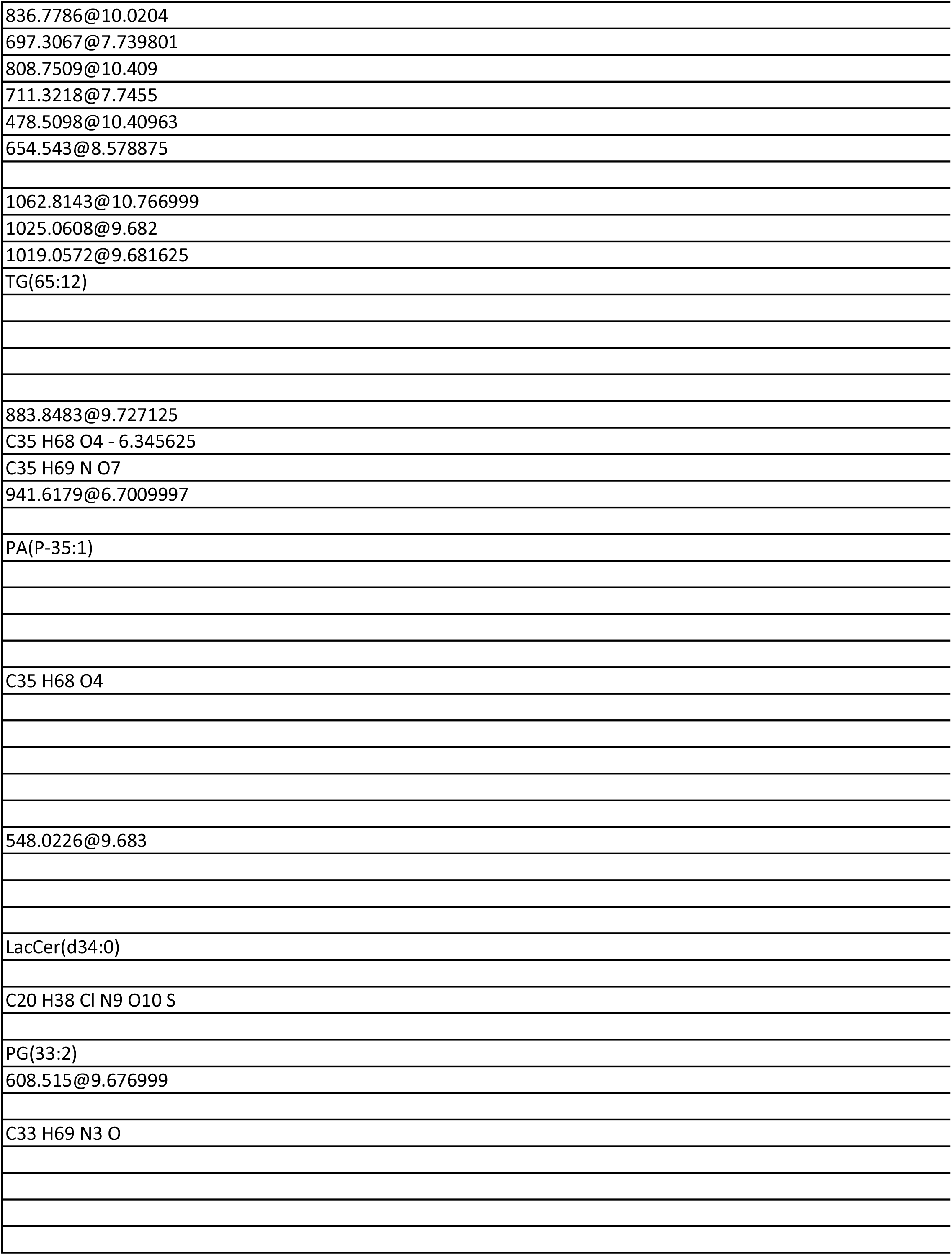
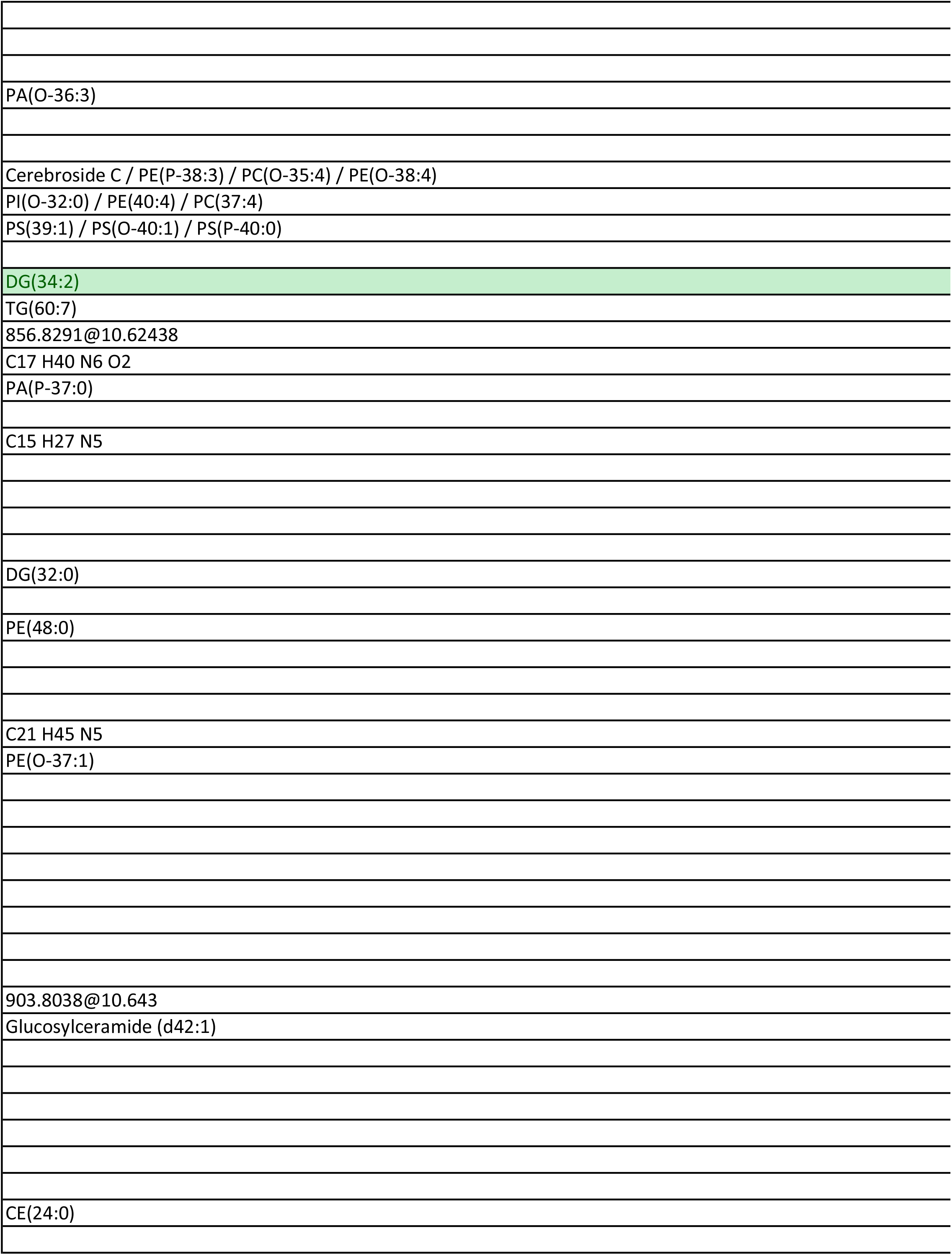
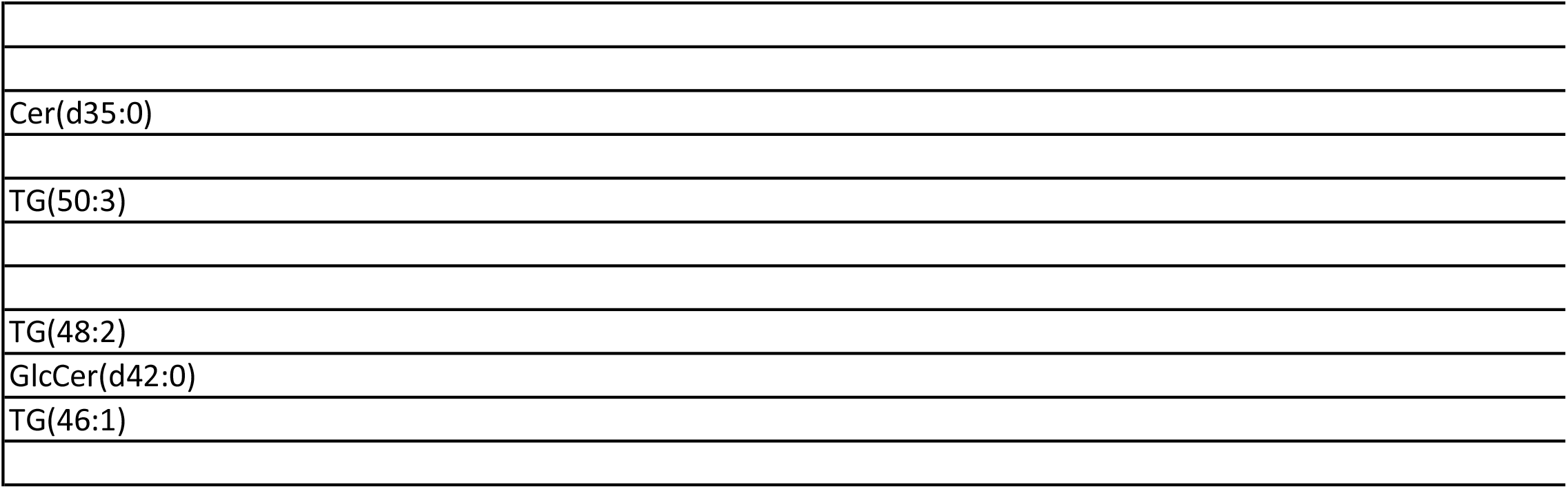

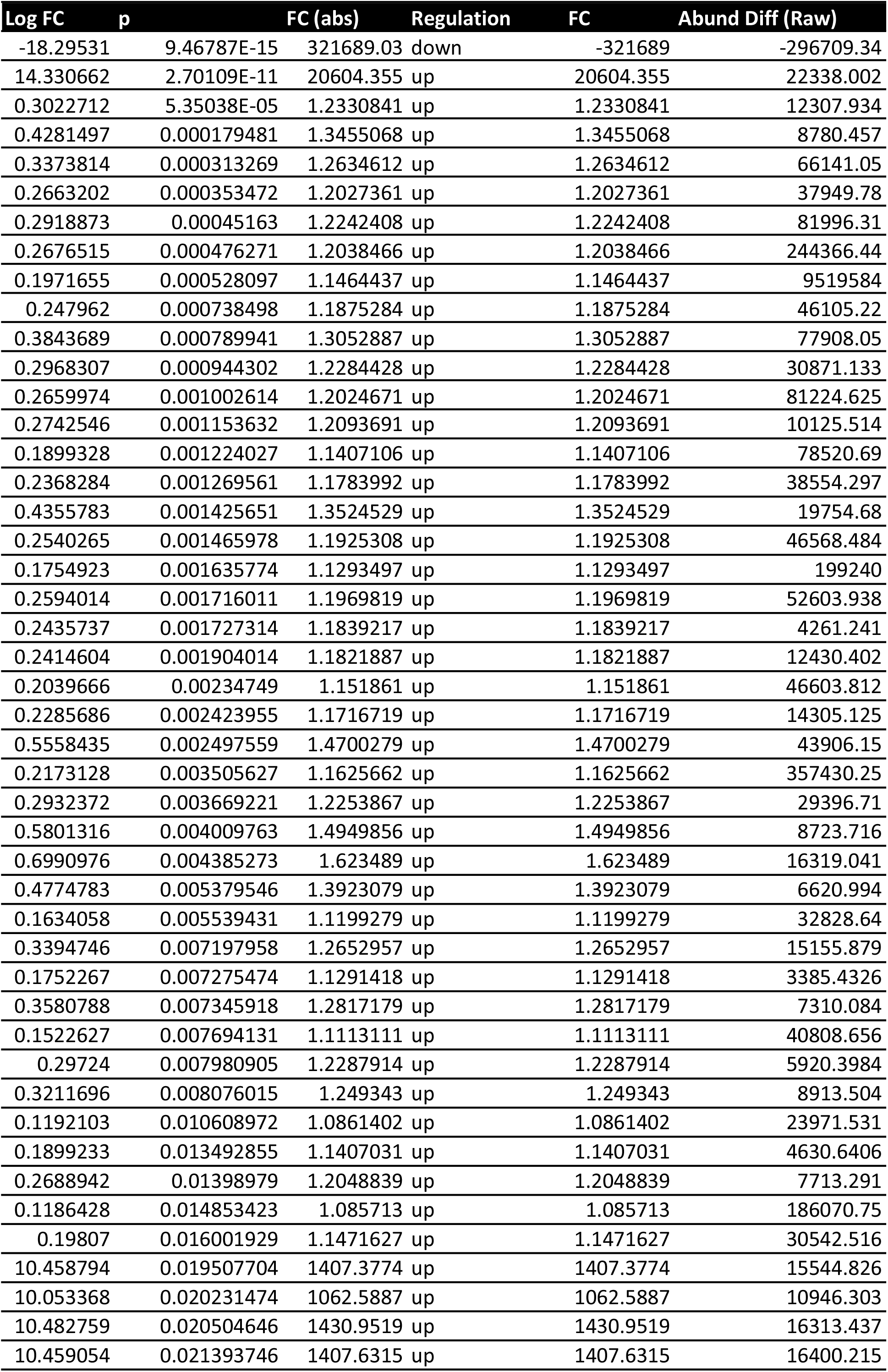
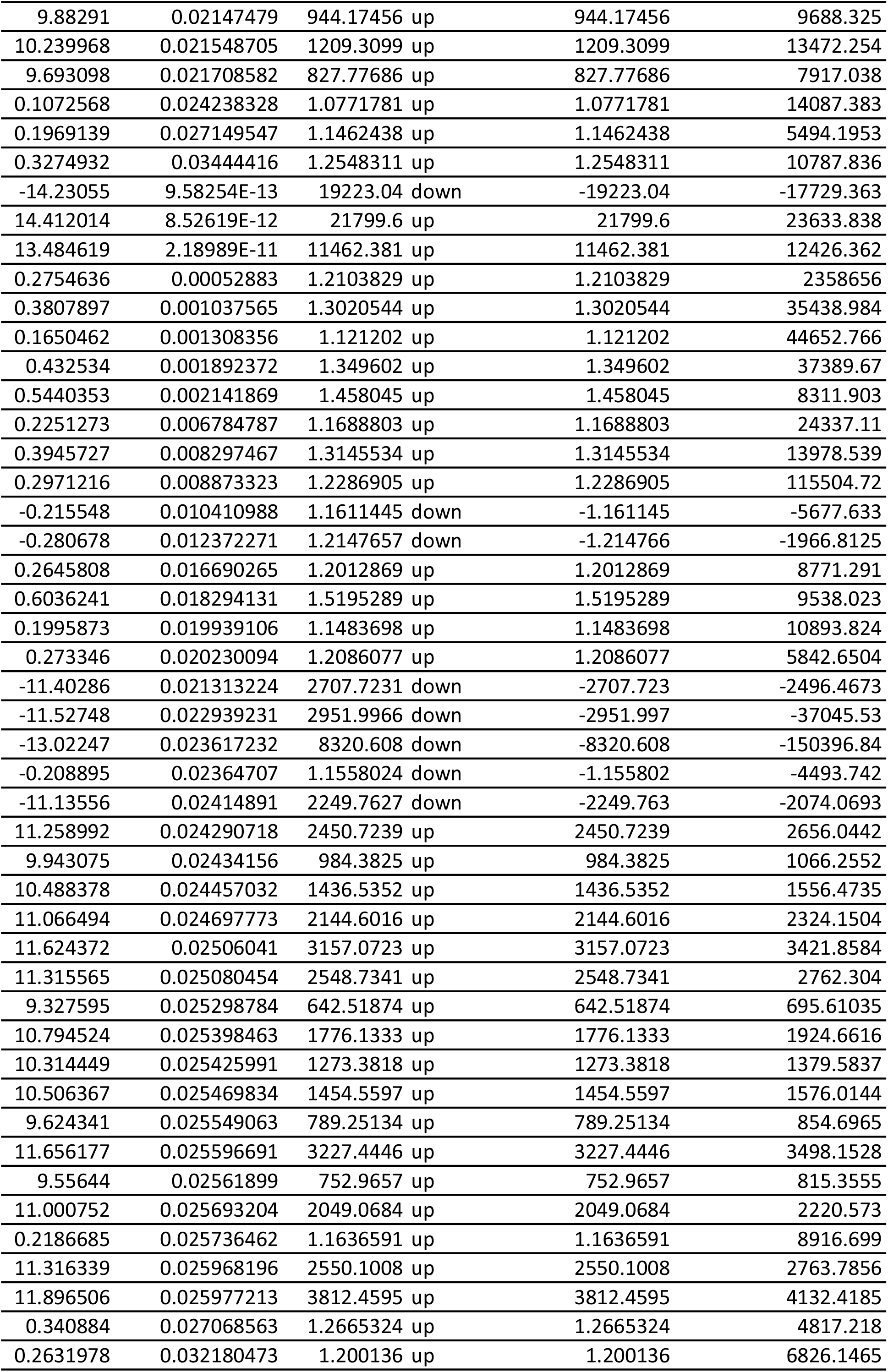
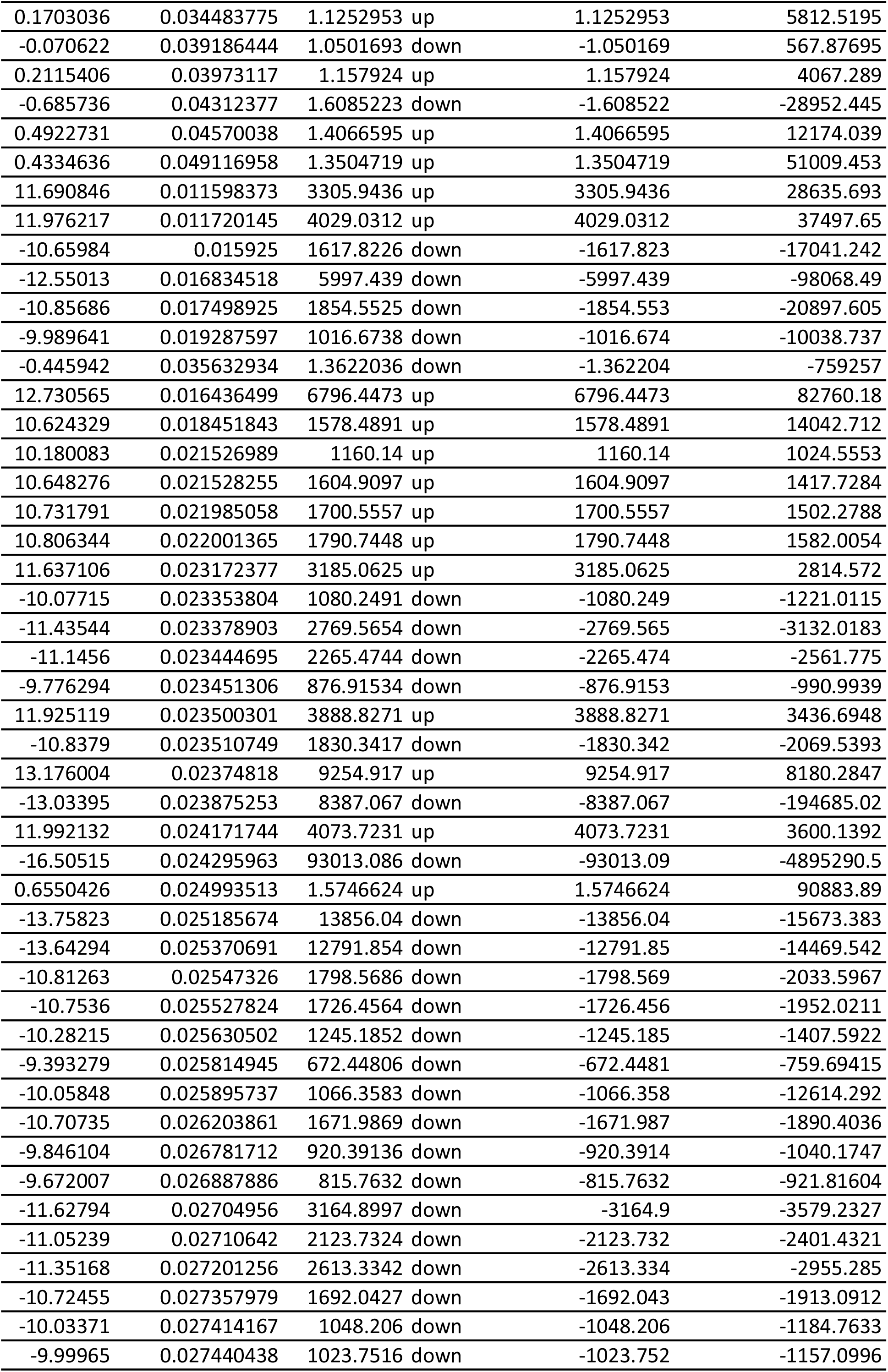
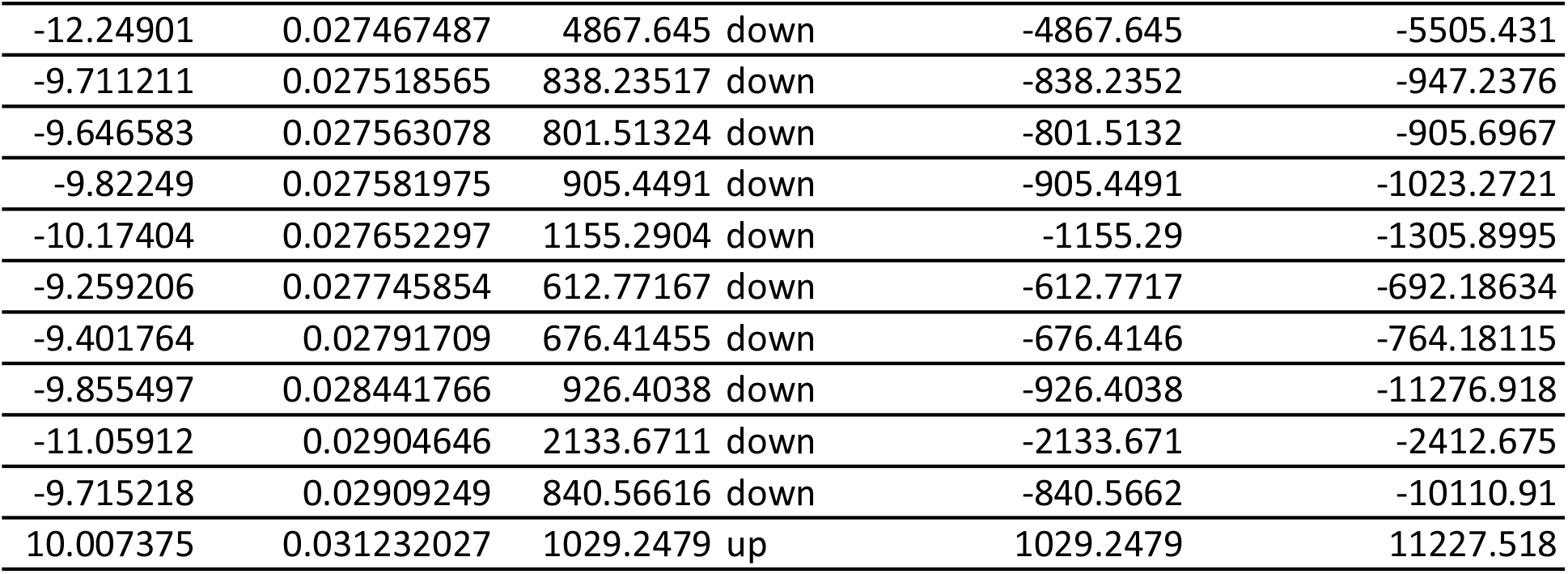

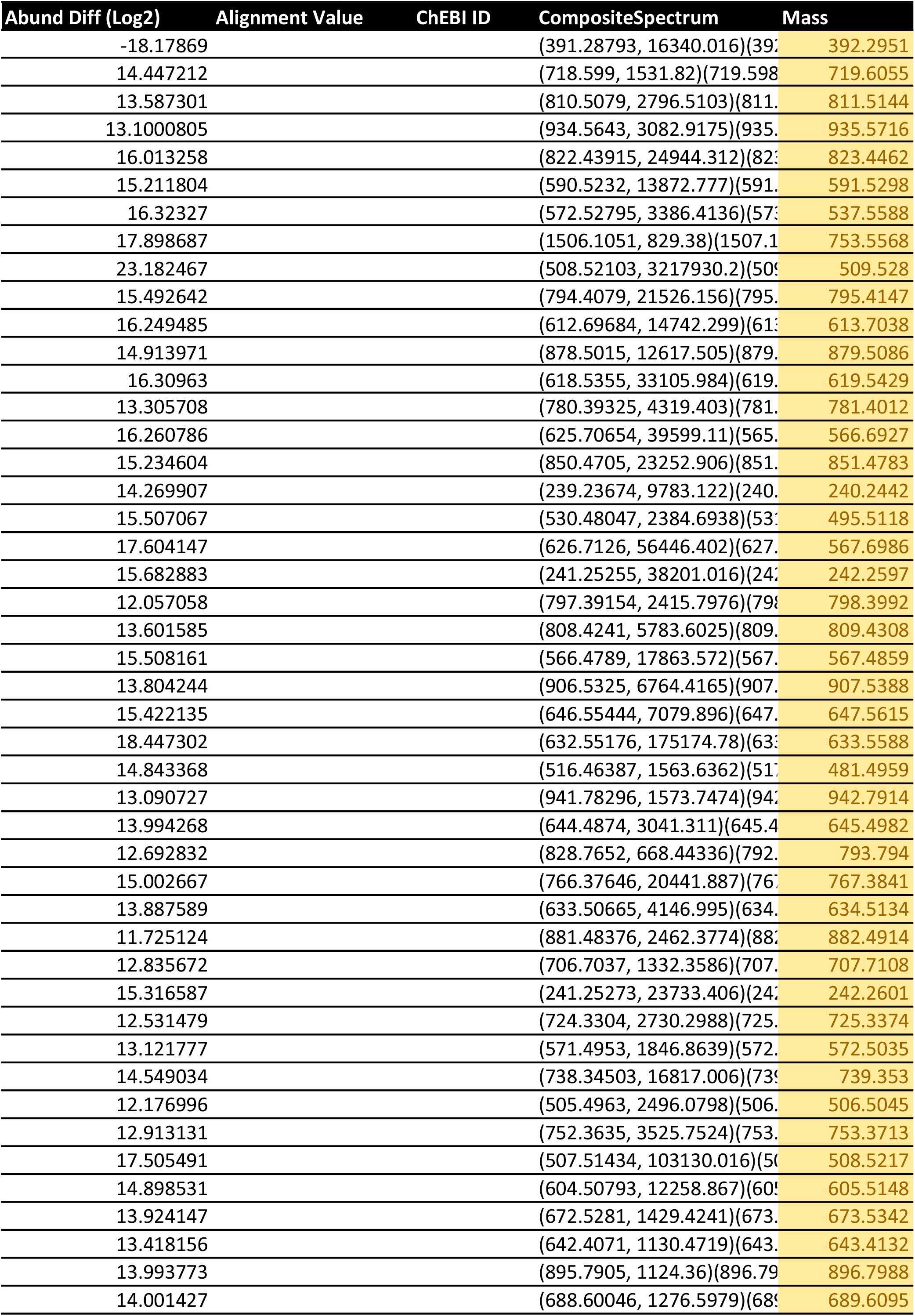
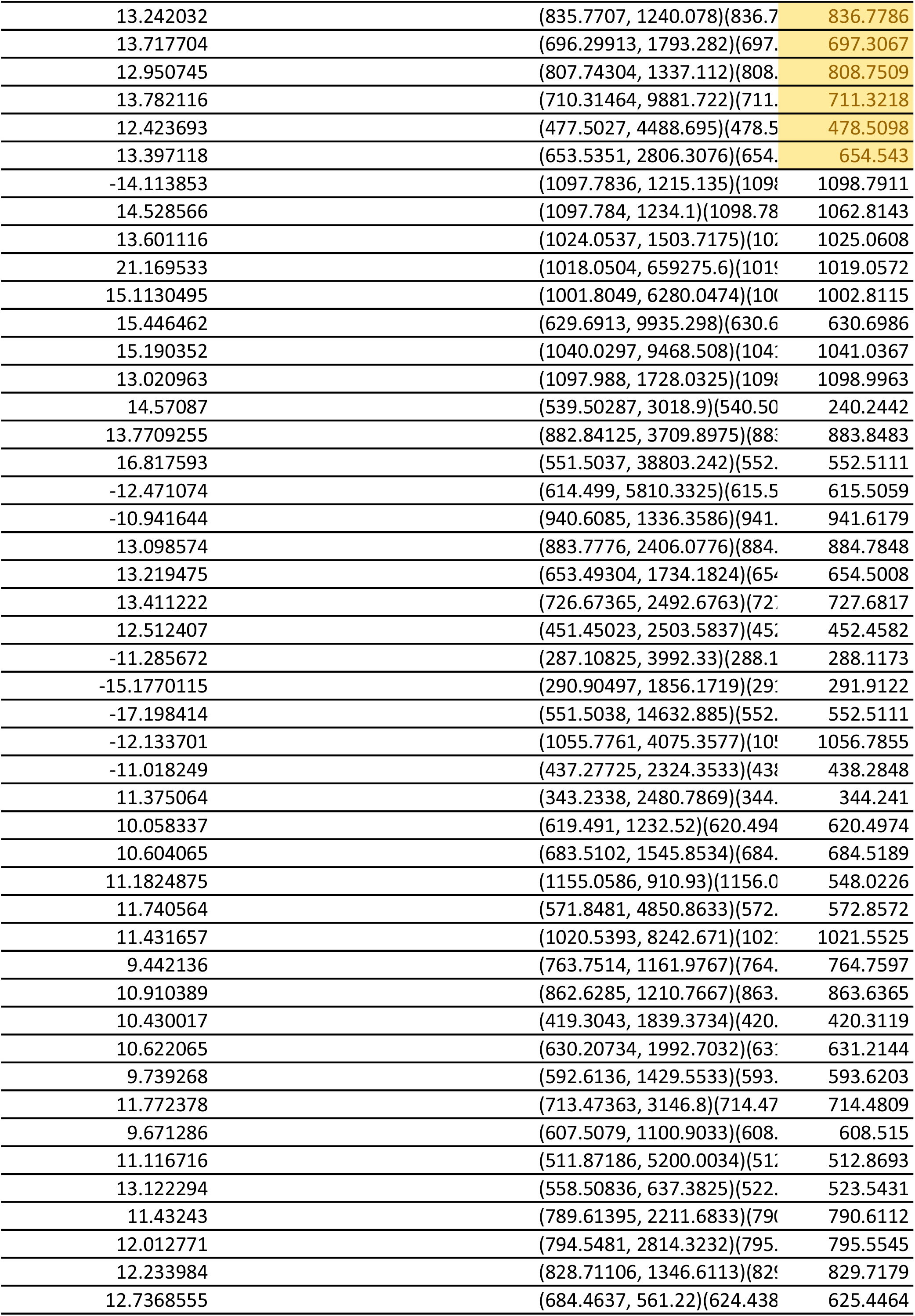
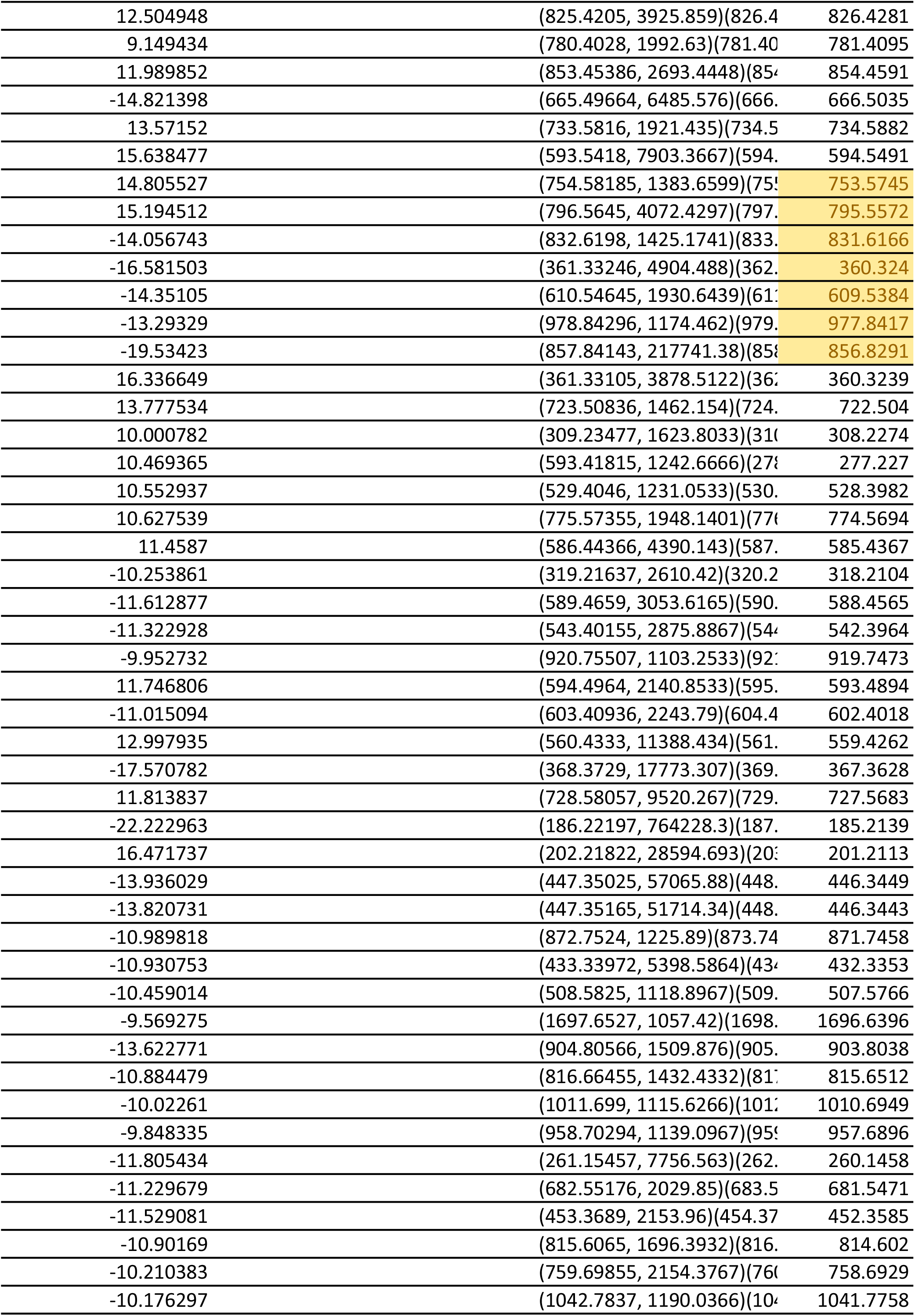
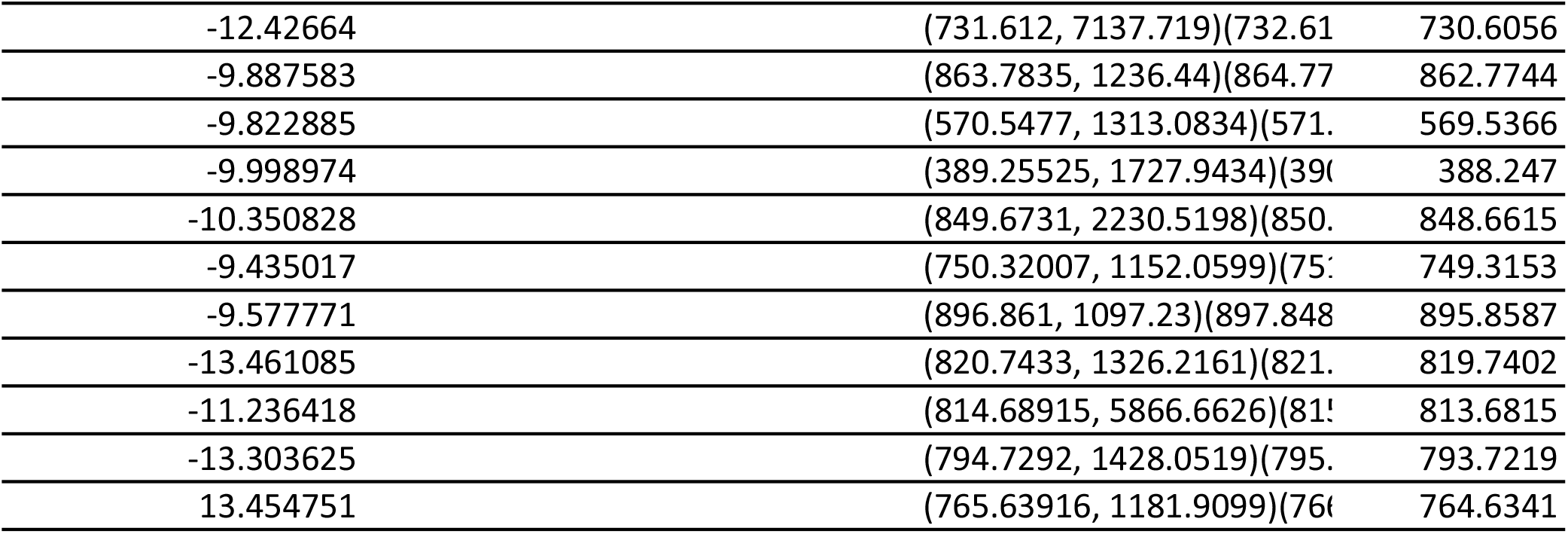

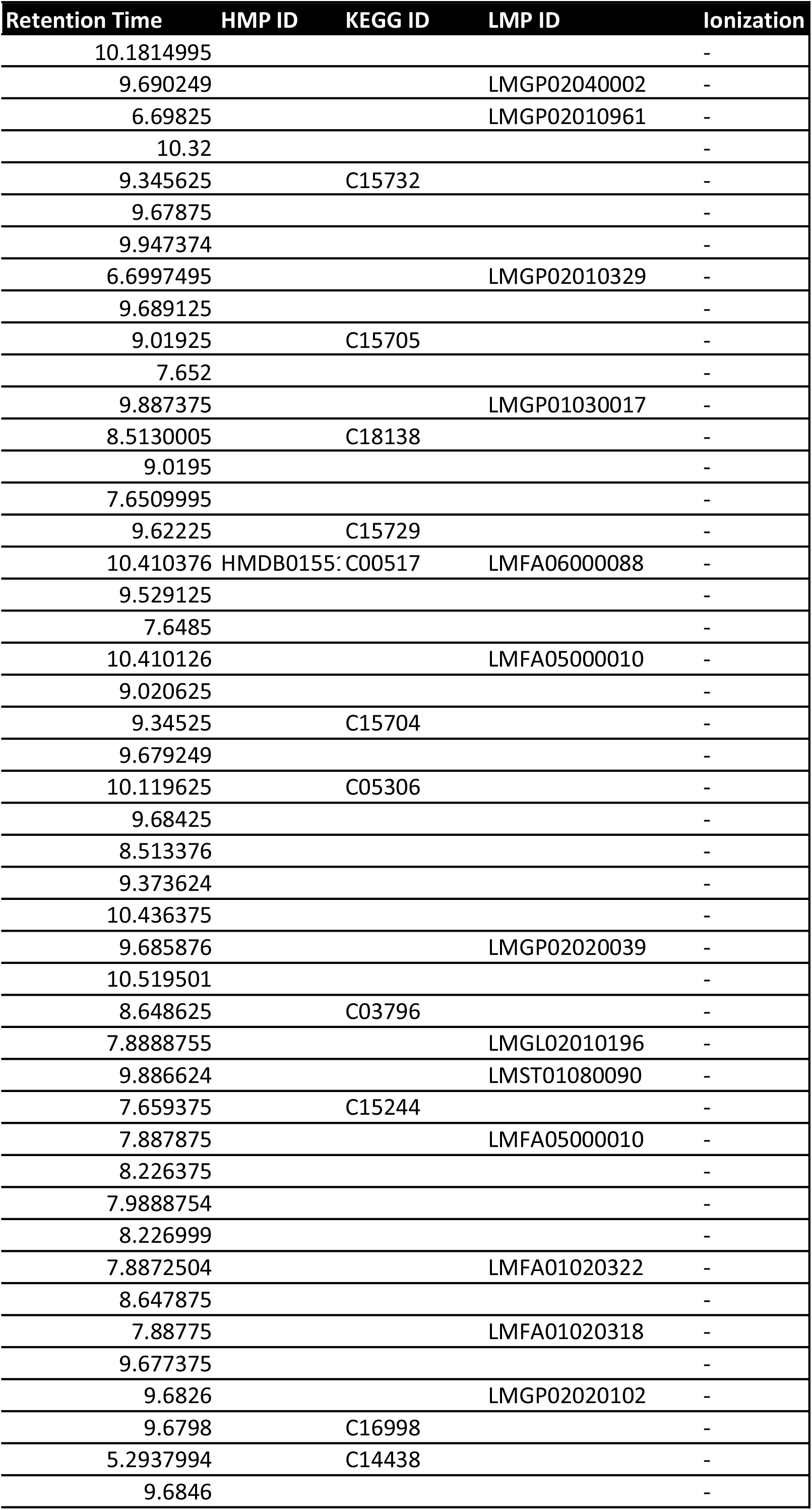
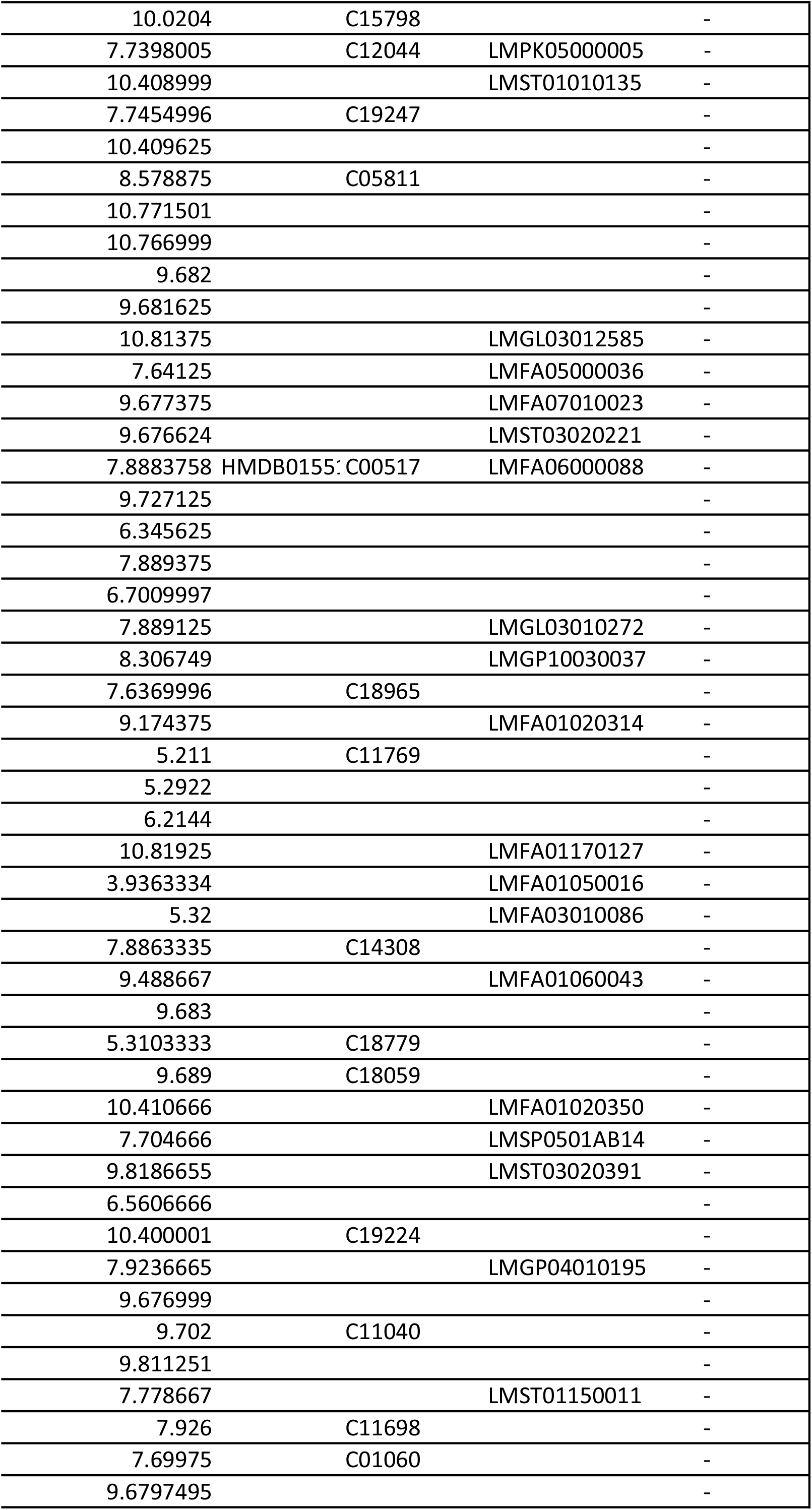
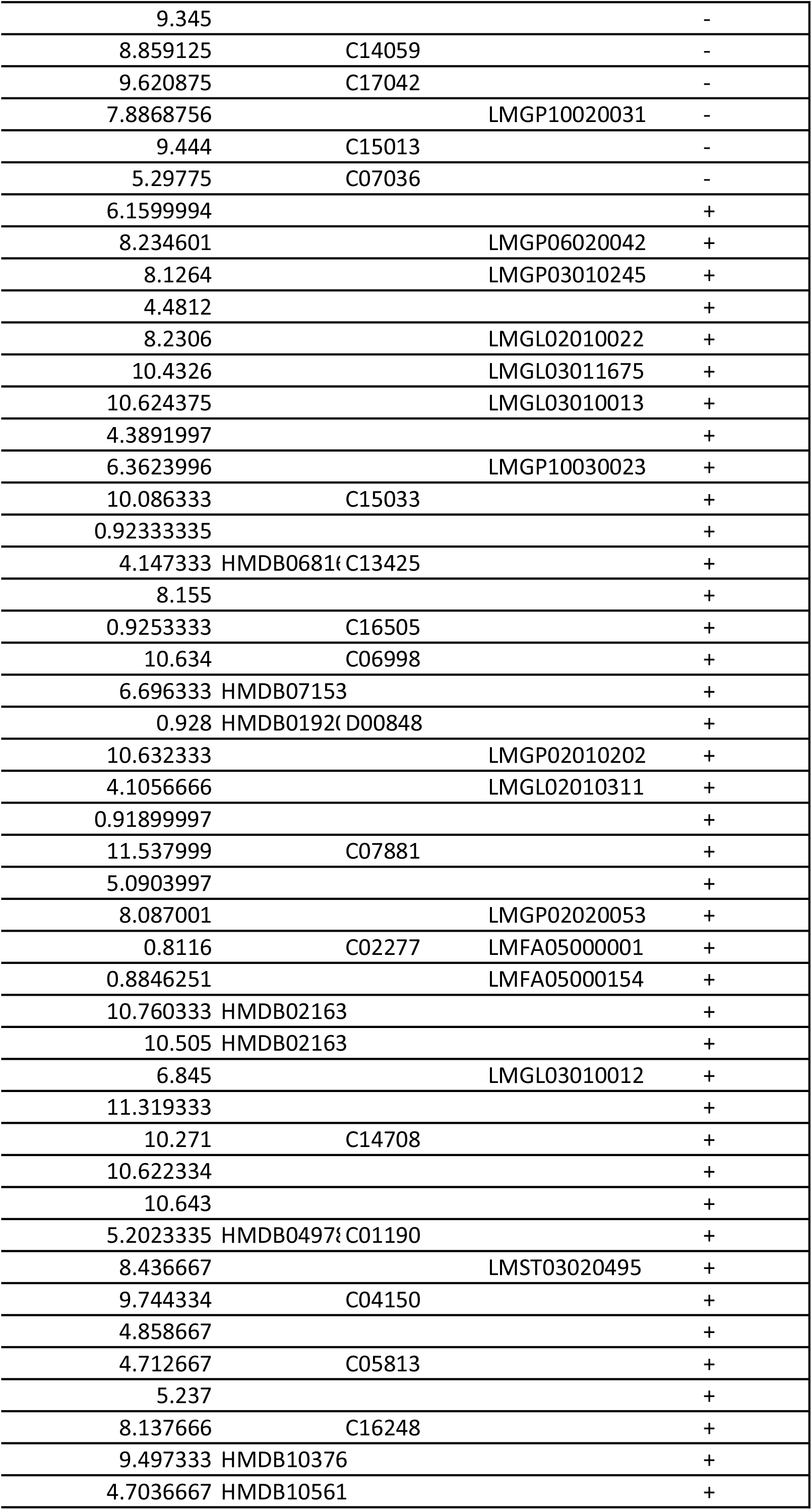
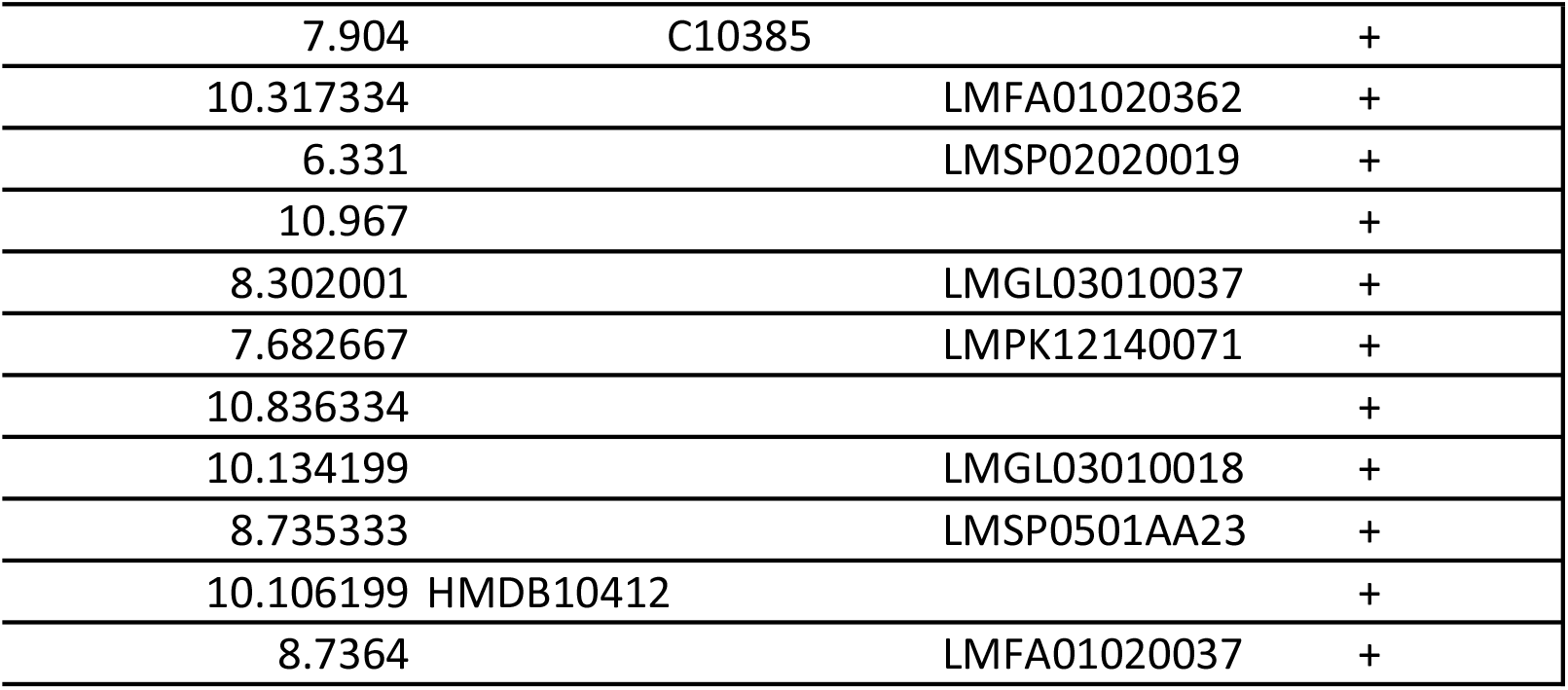

